# Novel YAP1/TAZ pathway inhibitors identified through phenotypic screening with potent anti-tumor activity via blockade of GGTase-I / Rho-GTPase signaling

**DOI:** 10.1101/2023.08.30.555331

**Authors:** Keith Graham, Philip Lienau, Benjamin Bader, Stefan Prechtl, Jan Naujoks, Ralf Lesche, Joerg Weiske, Julia Kuehnlenz, Krzysztof Brzezinka, Lisette Potze, Francesca Zanconato, Barbara Nicke, Anna Montebaur, Wilhelm Bone, Sven Golfier, Stefan Kaulfuss, Charlotte Kopitz, Sabine Pilari, Holger Steuber, Sikander Hayat, Atanas Kamburov, Andreas Steffen, Andreas Schlicker, Philipp Buchgraber, Nico Braeuer, Nuria Aiguabella Font, Tobias Heinrich, Lara Kuhnke, Katrin Nowak-Reppel, Carlo Stresemann, Patrick Steigemann, Annette O. Walter, Simona Blotta, Matthias Ocker, Ashley Lakner, Dominik Mumberg, Knut Eis, Stefano Piccolo, Martin Lange

## Abstract

This study describes the identification and target deconvolution of novel small molecule inhibitors of oncogenic YAP1/TAZ activity with potent anti-tumor activity in vivo. A high-throughput screen (HTS) of 3.8 million compounds was conducted using a cellular YAP1/TAZ reporter assay. Target deconvolution studies identified the geranylgeranyltransferase-I (GGTase-I) complex, as the direct target of YAP1/TAZ pathway inhibitors. The novel small molecule inhibitors block the activation of Rho-GTPases, leading to subsequent inactivation of YAP1/TAZ and inhibition of cancer cell proliferation in vitro. Multi-parameter optimization resulted in BAY-593, an in vivo probe with favorable PK properties, which demonstrated anti-tumor activity and blockade of YAP1/TAZ signaling *in vivo*.

**SIGNIFICANCE:** YAP1/TAZ have been shown to be aberrantly activated oncogenes in several human solid tumors, resulting in enhanced cell proliferation, metastasis and provision of a pro-tumorigenic microenvironment, making YAP1/TAZ targets for novel cancer therapies. Yet, the development of effective inhibitors of these potent oncogenes has been challenging. In this work, we break new ground in this direction through the identification of novel inhibitors of YAP1/TAZ activity.

**Graphical abstract:** 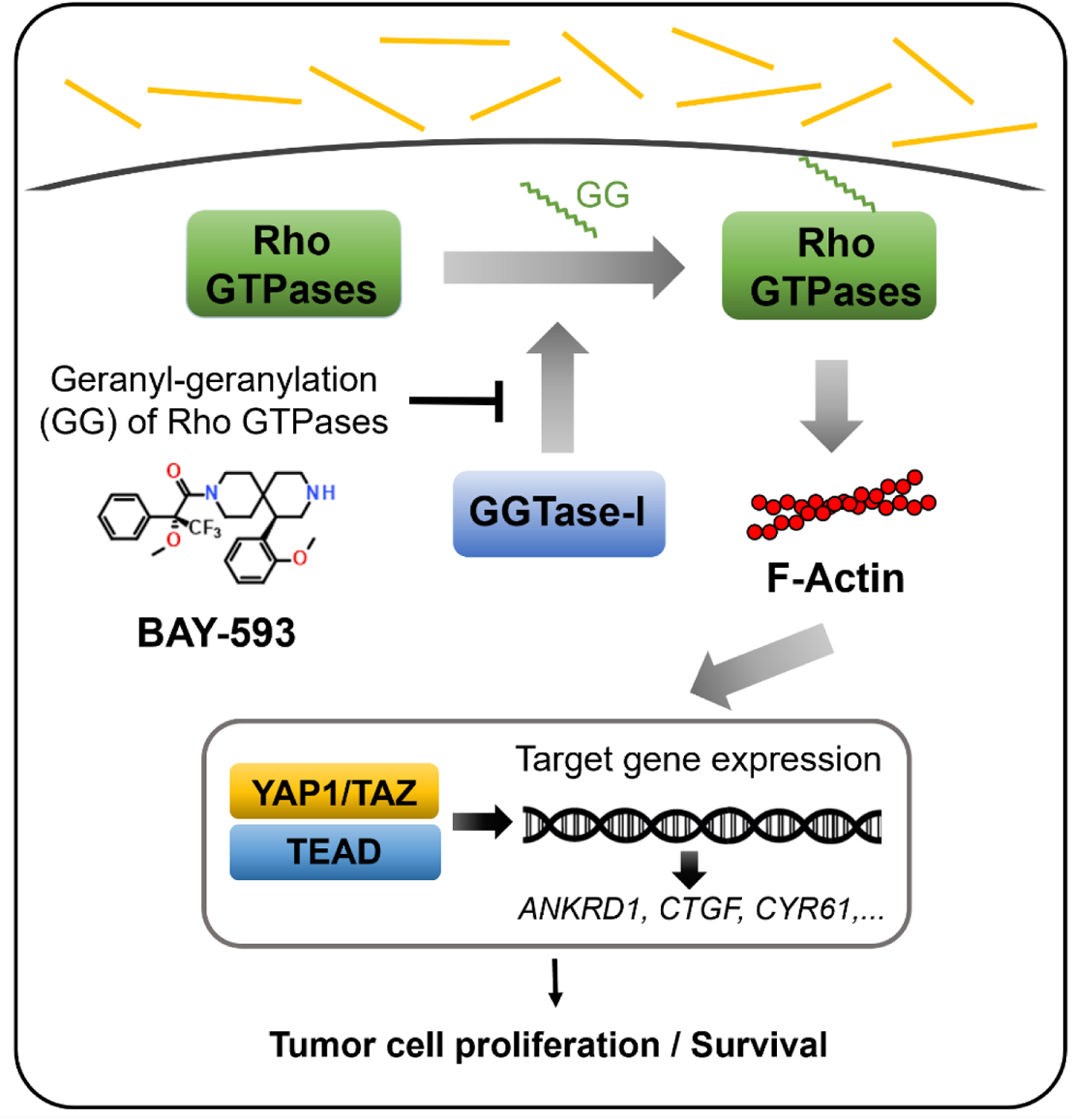

**HIGHLIGHTS:**

- Novel YAP1/TAZ pathway inhibitors identified by phenotypic high-throughput screen
- Target deconvolution identifies GGTase-I as the direct target of the novel YAP1/TAZ pathway inhibitors
- GGTase-I inhibitors block Rho-GTPase signaling and downstream YAP1/TAZ
- GGTase-I inhibitor BAY-593 demonstrates potent anti-tumor activity *in vivo*

## INTRODUCTION

Yes-associated protein (YAP1) and TAZ (transcriptional coactivator with PDZ-binding motif also known as WWTR1) protein are closely related transcriptional co-activators. YAP1/TAZ have been identified as downstream effectors of the Hippo pathway regulating organ size and cell proliferation during embryonic development and tissue repair^1–4^. Upstream components of the Hippo pathway consist of a set of serine/threonine kinases, such as sterile 20-like kinase 1/2 (MST1/2) and large tumor suppressor 1/2 (LATS1/2) which phosphorylate and inhibit the downstream effectors YAP1/TAZ, leading to cytoplasmic retention and degradation^5,6^. Frequently observed loss-of-function mutations in the Hippo pathway components and tumor suppressor neurofibromatosis 2 gene (*NF2*) lead to aberrant YAP1/TAZ activation in mesotheliomas, schwannomas and meningiomas ^7,8^. Active YAP1/TAZ form complexes with TEA domain (TEAD) family member transcription factors at promoters and enhancers in the nucleus, leading to activation of pro-tumorigenic target genes including *CTGF*, *CYR61, ANKRD1* and *AXL*, amongst others ^9,10^. The expression of target genes subsequently leads to promotion of cell proliferation and inhibition of apoptosis ^11^. Based on evidence from transgenic mouse models, aberrant activation of YAP1/TAZ is a hallmark of cancer resulting in enhanced cancer cell proliferation, metastasis and provision of a pro-tumorigenic microenvironment ^7,12–14^. These findings make YAP1/TAZ promising targets for anti-cancer therapy.

YAP1/TAZ are regulated by several signaling cascades including the Hippo, 5’ adenosine monophosphate-activated protein kinase (AMPK), Wnt, G protein-coupled receptor-regulated and mechanotransduction signaling pathways, offering various routes for therapeutic intervention^15^. YAP1/TAZ signaling is crucial for cancer and wound healing, however, it appears to be dispensable at least in some tissues for normal tissue homeostasis in adults ^16^.

Currently, no YAP1/TAZ inhibitors have been approved for cancer treatment. Several small molecule inhibitors and peptides targeting the Hippo pathway or the YAP1/TAZ transcriptional complex directly have been described in the literature ^17–19^. Recently, a small molecule targeting the YAP1/TAZ transcriptional co-activators TEAD has advanced to clinical trials ^20^. Here, we describe the identification, optimization and characterization of novel small molecule YAP1/TAZ pathway inhibitors.

## EXPERIMENTAL PROCEDURES (MATERIALS AND METHODS)

### Compounds

BAY-856 (hit compound), BAY-6092 (in vitro probe) and BAY-593 (in vivo probe) were synthesized at Bayer AG (Germany). The syntheses are described in WO2020/048830, Example 2-2; WO2020/048831, Example 89 and WO2020/048829 Example 62, respectively. RhoA inhibitors, C3 exoenzyme, cerivastatin, GGTI-298 and GGTI-2133, lonafarnib and latrunculin A were obtained from Tocris Bioscience. The selectivity of BAY-856 was assessed using a kinase panel (Eurofins Discovery) consisting of 337 kinases measured at concentrations of 10µM ATP and 1µM BAY-856.

### Cell lines

769P, 786O, A-2058, CAL27, CFPAC-1, CHL-1, HEL92.1.7, H1373, H358, HT-1080, JJN-3, MDA-MB-231, MOLM-16, SCC9, SKMEL-3, T24 and T98G cells were purchased either from the American Type Culture Collection (ATCC) or the German Collection of Microorganisms and Cell Cultures (Deutsche Sammlung von Mikroorganismen und Zellkulturen, DSMZ) and cultured according to supplier’s protocols at +37°C in a humidified atmosphere containing 5% CO_2_. Before use, cell lines were tested to be free from mycoplasma contamination and subjected to DNA fingerprinting for authentication at DSMZ. MDA-MB-231 TEAD-luc cells were cultured in DMEM low glucose, 10% FBS, 1% Glutamax, 250 μg/mL hygromycin and 0.5 μg/mL puromycin.

### YAP1/TAZ reporter assay

The YAP1/TAZ reporter assay utilized a dual readout approach using MDA-MB-231 cells with stable expression of Luc2P firefly luciferase reporter (pGL4.27 vector, Promega) under control of a TEAD promoter element destabilized by PEST-domains and thymidine kinase (TK)-Renilla reporter (pGL4.74 vector, Promega) for toxicity control. Firefly luminescence followed by the Renilla luminescence were measured using the DualGlo™ luciferase assay system detection kit (Promega, #E2920, #E2940). The assay was performed on white 384-well or 1536-well microplates (Greiner Bio-One) with a total volume of 5 µL or 4 µL, respectively. Using either a Hummingbird liquid handler (Digilab) or an Echo® Acoustic liquid handler (Labcyte), 50 nl (40 nL in 1536-well microplates) of a 100-fold concentrated solution of test compounds in DMSO were transferred into a 384-well microtiter test plate, and 5 µL of freshly prepared cell suspension was added to the wells, followed by incubation at +37°C for 20-24 hours. For dose-response evaluation, compounds were tested in duplicate at up to 11 concentrations by serial dilution (20 µM, 5.7 µM, 1.6 µM, 0.47 µM, 0.13 µM, 38 nM, 11 nM, 3.1 nM, 0.89 nM, 0.25 nM and 0.073 nM). After incubation, 1 µL of the Dual-Glo™ Luciferase detection solution was added to the wells, and the test plates were centrifuged and incubated at +20°C for 10 minutes before the luminescence signal was measured using either a Pherastar (BMG) or a ViewLux™ (PerkinElmer) microplate reader. Subsequently, Dual-Glo™ Stop&Glo Luciferase detection solution (1 µl/well) was added to the wells. The plates were centrifuged and incubated at +20°C for 10 minutes before measuring the TK-Renilla luminescence signal. Data were normalized to control wells containing cells without test compounds (0% inhibition) or with control inhibitor latrunculin A (100% inhibition).

### YAP1/TAZ translocation assay

The YAP1/TAZ translocation assay utilizing high-content analysis (HCA) technology was used for measuring the shuttling of YAP1/TAZ from the nucleus to the cytoplasm. MDA-MB-231 cells were seeded at a density of 2,000 cells/well into a 384-well microtiter plate in 10 μl medium and incubated at +37°C overnight. The cells were treated with BAY-856 at indicated concentrations (0.005 µM – 10 μM or DMSO for vehicle control, final volume of 20 μl) at +37°C for 24 hours, fixed with 4% paraformaldehyde (PFA) and permeabilized with 0.5% Triton™ X-100 in blocking solution containing 1% bovine serum albumin (BSA) in phosphate buffered saline (PBS). After washing with PBS, YAP (D8H1X) XP® Rabbit monoclonal antibody (Cell Signaling (CST), #14729) and Hoechst 33342 nucleic acid stain (Thermo Fisher Scientific, H3570) were used for labelling the phosphorylated YAP1/TAZ-complex and nuclei for HCA microscopy, respectively. High-throughput HCA microscopy images were acquired using a Perkin Elmer PHENIX™ reader and analyzed with MetaXpress™ image analysis software (Molecular Devices, version 6.2.3).

### Cellular thermal shift assay–mass spectrometry (CETSA^®^-MS)

Target engagement of BAY-856 and BAY-6092 was determined by cellular thermal shift assay combined with mass spectrometry (CETSA^®^-MS), Pelago Bioscience AB) in MDA-MB-231 cells. The studies were performed according to the general CETSA^®^ protocol ^21^ and the optimized CETSA MS protocol ^22^. Briefly, MDA-MB-231 cell lysates were treated with 10 μM BAY-856, BAY-6092 or DMSO control and incubated under constant agitation at room temperature (RT) for 30 minutes. The lysates were divided into 60 μl aliquots and subjected to a 10-step heat exposure between +40-67°C for 3 minutes. Precipitated proteins were centrifuged at 20,000 × g for 20 minutes and supernatants containing the soluble fractions were used for further MS analyses. After the reduction, denaturation and digestion steps, peptide fractions were labelled with 10-plex Tandem Mass Tag (TMT10) reagents (Pierce) at RT for 4 hours. The labelled samples were subsequently acidified and desalted using a spin cation exchanger (Strata X, Phenomenex), eluted in acetonitrile:water (60:40) containing 0.5% acetic acid and dried using a centrifugal evaporator. The samples were dissolved in 20 mM ammonia (pH 10.9) and subjected to reversed-phase high pH fractionation on an ÄKTA Micro HPLC system (GE Healthcare) using a 2.1 × 250 mm C18 column (Xbridge Peptide BEH C18, 300 Å, 3.5μm particle size, Waters Corporation). The collected fractions were pooled, evaporated to dryness and analyzed by high resolution nanoscale liquid chromatography tandem mass spectrometry (nanoLC-MS/MS) on Q-Exactive Orbitrap mass spectrometers (Thermo Fischer Scientific) coupled to high performance nano LC systems (Dionex UltiMate 3000, Thermo Fischer Scientific). MS/MS data were collected on the Q Exactive using higher energy collisional dissociation (HCD). Protein identification was performed using the Sequest HT algorithm as implemented in the ProteomeDiscoverer 2.1 software package.

### Thermal shift assay (TSA)

TSA experiments were carried out in a 384-well plate format with 5 µL reaction volume using a recombinant PGGT1B and a ViiA™ 7 Real-Time PCR system (Thermo Fisher Scientific). Melting curves for recombinant PGGT1B protein were obtained at a protein concentration of 1.0 µM and 6xSYPRO Orange (Invitrogen) using a buffer containing 50 mM HEPES pH 7.5, 40 mM NaCl, 5 µM ZnCl_2_ and 1.0 mM TCEP. For binding experiments, the hit compound BAY-856 was added from 10 mM stock solution to a final concentration of 100 µM. Control samples contained 1% DMSO. Scans were measured from +25°C to +95°C at a scanning rate of +4°C/min. All TSA data were analyzed using Genedata Assay Analyzer software (Genedata, version 13.0).

### CRISPR screen

The MDA-MB-231 TEAD-luc Cas9 cell line was generated by lentiviral delivery of Cas9 in pLv5-Cas9-Neo (Sigma-Aldrich) into MDA-MB-231 TEAD-luc reporter cells, followed by antibiotic selection with Geneticin (Sigma-Aldrich) and clonal expansion. The derived clonal cell lines were validated for Cas9 activity and TEAD-luc reporter activity using a CRISPRuTest™ Functional Cas9 Activity Assay Kit (Cellecta) and Dual-Glo Luciferase Assay System (Promega), respectively. To perform a large-scale pooled CRISPR screen, the assay readout was established to detect the actual luciferase protein expression induced by TEAD-promoter activation. Luciferase protein expression was determined by flow cytometry. The cells were fixed, permeabilized and stained for firefly luciferase expression using a rabbit monoclonal anti-firefly luciferase antibody (Abcam, ab185924), followed by staining with a goat anti-rabbit Alexa Fluor 488 secondary antibody (Thermo Fisher Scientific, A-11008).

For large scale and focused custom single guide RNA (sgRNA) library screening, cells were transduced with Module 2 of the 150k 3-Module pooled Human Genome-Wide CRISPR Library (Cellecta) targeting 6,000 human protein-coding genes with up to 8 sgRNAs per gene or two custom libraries targeting 81 genes (Cellecta 10 sgRNAs/target and 10 shRNAs/target, respectively). A custom sgRNA (Cellecta) targeting the firefly luciferase reporter gene was spiked into Module 2 of the 150k 3-Module pooled Human Genome-Wide CRISPR Library as positive control. 50-100 × 10^6^ cells per module were transduced at a multiplicity of infection (MOI) of 0.5 (500-1000-fold library coverage). After puromycin selection and cell expansion, the resulting pool of gene-edited cells was split in two populations, re-plated at low density and treated with 37.4 nM BAY-856 (IC_20_) or left untreated for 24 hours. On day 15 post transduction, cells were fixed using the Fixation/Permeabilization Solution Kit (BD Biosciences) and stained for firefly luciferase expression (TEAD-luc reporter) using an anti-firefly luciferase antibody (Abcam, ab185924) followed by goat anti-rabbit Alexa Fluor 488 secondary antibody staining. Single cells were sorted on 10% firefly luciferase low and 10% firefly luciferase high expression using fluorescence-activated cell sorting (FACS). At least 50 × 10^6^ cells (500-fold library coverage) were collected for the luciferase low and high bin, respectively. Genomic DNA was isolated using the GeneRead DNA FFPE Kit (Qiagen). The Illumina sequencing libraries were generated via two-step PCR amplification using primer sequences provided by Cellecta, summarized in **Supplementary Table 1**. The sgRNA abundance in sorted and treated cells was sequenced on a HiSeq 2500 sequencing system (Illumina) with a sequencing depth of at least 1,000 reads per sgRNA. Sequencing reads were aligned to the respective library using Bowtie short read aligner open-source software (version 2.2.6) with no mismatches allowed. Generated read counts were normalized for each sample and fold change (FC) between the luciferase low and high fractions was calculated for individual sgRNAs within each module. To determine the compound effect, firefly luciferase low-high-fold-changes between BAY 856- and untreated cells were plotted against each other and compared. For single knockouts, MDA-MB-231 TEAD-luc Cas9 cells were transduced with sgRNAs against the *PGGT1B* gene. Four days after transduction, the cells were re-seeded in respective formats, treated with BAY-856, and TEAD-luc reporter assay was performed.

### Biochemical geranylgeranyl assay

BAY-856-mediated inhibition of the enzymatic activity of purified human GGTase-I was measured using a biochemical GGTase-I assay. The assay was established according to Mansha *et al.* (Mansha et al., 2016) and further optimized and miniaturized to result in the high-throughput amenable add-only assay used in this study. The β subunit of human GGTase-I (PGGT1b, amino acids M1-T377) and the α subunit of human farnesyltransferase (FNTA, amino acids M1-Q379), which are essential constitutes of the functional GGTase-I complex, were expressed in Hi-5 insect cells and purified by size exclusion chromatography. The assay was performed using white 384-well microplates (Greiner Bio-One) with a total volume of 5 µl/well. BAY-856 (50 nL,100-fold concentrated solution in DMSO) was transferred into 384-well plates using a Hummingbird liquid handler, and 2.5 µL of 5 nM GGTase-I in assay buffer (50 mM Tris-HCl pH 7.4, 5 mM MgCl_2_, 10 mM KCl, 50 µM ZnCl_2_, 5 mM DTT, 0.04% n-dodecyl-β-D-maltoside, 5 mM ATP) was added to the wells. After pre-incubation at RT for 15 minutes, the reaction was started by adding 2.5 µL of 2 µM dansyl-GCVLL peptide and 2 µM GGPP in assay buffer. The plates were incubated for 60 minutes at RT, and the fluorescence intensity of the reaction mixture was measured with a Pherastar microplate reader at 380/510 nm. The fluorescence signal of BAY-856-treated samples was normalized to DMSO control (0% inhibition) containing complete reaction. The control sample (100% inhibition) contained complete reaction without either enzyme or BAY-856. For dose-response evaluation, BAY-856 was tested in duplicate at up to 11 concentrations (20 µM, 5.7 µM, 1.6 µM, 0.47 µM, 0.13 µM, 38 nM, 11 nM, 3.1 nM, 0.89 nM, 0.25 nM and 0.073 nM). Dilution series were prepared before the assay in a 100-fold concentrated form by serial dilution with two separate dilutions (n=2).

### X-ray crystallography

Purified rat GGTase-I (15 mg/mL) containing α and β subunits was incubated with the in vitro probe BAY-6092 and co-crystallized using the hanging drop vapor diffusion method by mixing 1 µL of the protein solution with 1 µL of reservoir solution containing 1.2 M K-Na-tartrate, 100 mM (NH_4_)_2_SO_4_, 100 mM MES pH 6.5 at +20°C. Cuboid-shaped crystals of approximately 100 µm of length were harvested from the crystallization drop, cryoprotected using 25% 2-methyl-2,4-pentanediol (MPD) in the reservoir solution and flash-frozen in liquid nitrogen. The X-ray diffraction data for the GGTase-I/BAY-6092 complex was collected at PETRA (DESY, Hamburg, Germany) and processed using the XDS program package. The structure was solved by molecular replacement using PHASER (Mccoy et al., 2007) with PDB entry 1N4P (Taylor et al., 2003) as a search template. Model building and refinement were carried out using COOT ^23^ and REFMAC ^24^, respectively.

### Cell fractionation and protein detection by capillary electrophoresis

Cell fractionation was performed using a Subcellular Protein Fractionation kit (Thermo Fischer Scientific, #7884) and the protein levels were measured by capillary electrophoresis (Peggy Sue™, Protein Simple) according to manufacturer’s instructions. The antibodies used for protein detection are listed in the **Key Resources Table**.

### Protein immunoblotting

MDA-MB-231 whole cell lysates were obtained by sonication in lysis buffer containing 10 mM HEPES (pH 7.8), 150 mM NaCl, 5% glycerol, 0.5% NP-40, 5 mM EDTA, 1 mM dithiothreitol (DTT) and phosphatase and protease inhibitors. Samples were run on 4-12% NuPAGE-MOPS acrylamide gels (Thermo Fisher Scientific) and transferred onto polyvinylidene difluoride (PVDF) membranes by wet electrophoretic transfer. Blots were blocked with 0.5% non-fat dry milk and incubated with primary antibodies for GAPDH (Millipore, MAB374) and GFP FL (Santa Cruz Biotechnology, sc-8334) at +4°C overnight. Secondary antibodies anti-mouse HRP (Abcam, ab97023) and anti-rabbit HRP (Abcam, ab205718) were incubated for 1 hour at RT and then blots were developed using Pierce™ ECL Plus Western Blotting Substrate (Pierce, #32132). Images were acquired with ImageQuant LAS 4000 (GE Healthcare Life Sciences).

### RNA extraction and quantitative PCR

Quantitative PCR (qPCR) analyses were used to evaluate YAP1/TAZ target gene inhibition in MDA-MB-231 cells *in vitro* and in HT-1080 tumors *in vivo*. 1 × 10^5^ of MDA-MB-231 cells were plated in 500 µL/well in 24-well cell culture plates and incubated at +37°C overnight. Subsequently, BAY-856 (0.03 µM-10 µM) or vehicle (DMSO) were added using a D300e digital dispenser (Hewlett-Packard). The cells were incubated for 24 hours and lysed in 350 µL RLT buffer (Qiagen) prior to RNA extraction.

HT-1080 tumors were cut into small pieces and lysed in 300 µL RLT Plus buffer (Qiagen) with 40 µL of 1 M DDT using a TissueLyser II homogenizer (Qiagen) with stainless steel beads for three minutes. The lysates were centrifuged at maximum speed for three minutes, and supernatants were transferred into the gDNA Eliminator spin columns. Supernatants were centrifuged at 8,000 × g for 30 seconds and the flow-through was used for RNA extraction.

Total RNA was extracted using the RNAeasy Kit (Qiagen, #74181), and cDNA was prepared using the LunaScript® RT SuperMix Kit (New England Biolabs, #E3010) according to manufacturer’s instructions. qPCR reactions were performed using a Taqman qPCR system, Taqman Fast Advanced Master Mix (Thermo Fisher Scientific, #4444965) and Taqman Probes, listed in the Key Resources Table.

### RNA sequencing and bioinformatic analysis

TruSeq Stranded mRNA Library Prep (Illumina, RS-122-2103) was used to construct cDNA libraries starting with an input of 400 ng of total RNA per sample. cDNA libraries were pooled and clustered to a single-read sequencing flow cell (Illumina cBOT, HiSeq SR Cluster Kit v4, GD-401-4001 and sequenced on an Illumina HiSeq 2500 instrument (HiSeq SBS Kit v4 (50 Cycle Kit), FC-401-4002). On average, 19 million reads per sample were sequenced. Sample quality of the raw data (FASTQ) was checked through the internal NGS QC pipeline. The reads were mapped to the human reference genome (assembly hg19) using STAR aligner alignment tool (version 2.4.2a). FeatureCounts (version 1.4.6-p2) from the Subread package was used to compute the total read counts per gene based on the gene annotation downloaded from Gencode (version 19). The Bioconductor package DESeq2 was used for the differential gene expression analysis (DGEA) and the computation of FPKM expression values. Gene set enrichments analysis (GSEA) was conducted using the pre-ranked feature of the GSEA application (version 2.2) based on the ranking from the DGEA and MsigDB subsets C2 (curated gene sets), C3 (motif gene sets), C6 (oncogenic gene sets) as well as the ‘YAP1/TAZ/TEAD target gene’ gene set published by Zanconato et al.^25^. Sequencing raw data have been uploaded to the Gene Expression Omnibus (GEO) public repository (accession number GSE243122).

### OncoPanel™ Cell Proliferation screen

The OncoPanel™ Cell Proliferation (Eurofins) screen with 250 human cancer cell lines was performed to evaluate the anti-proliferative activity of the in vitro probe BAY-6092. Briefly, the cells were plated onto 384-well microplates and incubated at +37°C for 24 hours. Next, test compounds were added to the plates by a serial dilution in half-log steps of ten concentrations, ranging from 1 nM to 30 µM. All measurements were performed in duplicate. Control samples were treated with DMSO vehicle (0.1% final concentration). After 72 hours of treatment, cells were fixed and stained with DAPI for fluorescence imaging of nuclei. In order to determine cell proliferation at the end of treatment, a non-treated plate was prepared to serve as baseline before the initiation of treatments. Automated fluorescence microscopy was performed using an ImageXpress Micro XL high-content imager (Molecular Devices) with a 4X objective. Acquired images were analyzed by MetaXpress™ image analysis software (Molecular Devices, version 5.1.0.41). Cell proliferation was determined based on the fluorescence intensity of the incorporated nuclear dye. For each cell line, IC_50_, EC_50_ and activity area values for cell count were determined. Activity area is an estimate of the integrated area above the curve, ranging from 0-10, where 0 indicates no inhibition of proliferation at all concentrations, and 10 indicates complete inhibition of proliferation at all concentrations. Curve-fitting, calculations, and report generation were performed using a custom data reduction engine and MathIQ based software (AIM).

### Cell viability assay

The cells were plated onto 384-well microplates at a density of 300-1500 cells/well depending on their proliferation rate and incubated at +37°C for 24 hours. Next, test compounds were added to the plates using a D300 digital dispenser by 10-step 2,5-fold dilution series with final drug concentrations ranging from 0.1 nM to 100 nM. All measurements were performed in duplicate. Control samples were treated with DMSO vehicle (0.5% final concentration). After 72 hours of treatment, 30 µL of CellTiter-Glo (CTG) solution (Promega, #G755B) was added to each well. The plates were incubated at RT for 10 min, and the luminescence was measured using a VICTOR V multilabel plate reader (Perkin Elmer). In order to determine cell viability at the end of treatment, a non-treated plate was measured as baseline before the initiation of treatments. Relative number of viable cells and the half-maximal inhibition of cell viability (IC_50_) were calculated using the baseline plate (maximum inhibition) and the DMSO control (minimum inhibition). The IC_50_ values were calculated using a 4-parameter fit.

### Geranylgeranyl reporter assay

The capacity of the hit compound BAY-856 and the in vitro probe BAY-6092 to inhibit YAP1/TAZ downstream of GFP-RhoA or GFP-RhoA-F was assessed using MDA-MB-231 cells expressing the TEAD reporter (8xGTIIC-lux)^26^ and the TK-Renilla normalizer co-transfected with either GFP-RhoA with a geranylgeranylation motif or with a modified GFP-RhoA-F bearing a consensus motif for farnesylation as described ^27^. The cells were treated with 150 nM BAY-856, 2 nM BAY-6092, 1 μM GGTI-298, 0.1 μg/mL C3, 1 μM cerivastatin or vehicle (DMSO) at +37°C for 24 hours. The dual luciferase reporter readout was performed similarly to the YAP1/TAZ reporter assay described earlier. Results are expressed as mean±SD determined from three individual experiments performed in duplicate.

### Animals

All animal experiments were conducted in accordance with the German Animal Welfare Act and approved by local authorities. For MDA-MB-231 and HT1080 xenograft studies, 5-6 weeks old female NMRI nu/nu mice (Janvier) and nu/nu mice (Charles River) were used, respectively. The PXF 541 PDX model was performed at Charles River Discovery Research Services Germany GmbH using NMRI nu/nu mice. Mice were housed in pathogen-free and controlled conditions in individually ventilated cages and fed an irradiated soy-free diet and autoclaved tap water *ad libitum*.

### *In vivo* MDA-MB-231 TEAD-luc reporter study

1 × 10^6^ of MDA-MB-231 TEAD-luc cells suspended in 50% Matrigel / 50% medium were inoculated subcutaneously into the inguinal region of female mice. After 4 weeks, when tumors reached a volume of >100 mm^3^, animals were randomized (8-9 mice / group) and treated orally (per os, p.o.) with vehicle (90% PEG400, 10% water) or the hit compound BAY-856 twice daily (125 mg/kg; BID, twice daily) for 3 days. Bioluminescence imaging was performed 4 hours after drug administration on treatment days 1-3 using a NightOWL LB 983 In Vivo Imaging System (Berthold Technologies).

### *In vivo* efficacy studies

3 × 10^6^ HT-1080 cells or 5 × 10^6^ MDA-MB-231 cells suspended in 0.1 mL of 50% Matrigel / 50% medium were inoculated into the inguinal region of female immunocompromised mice. PXF 541 tumor pieces (approx. 4-5mm length) were implanted subcutaneously into the left flank of female immunocompromised mice. When tumors reached an average volume of 70-90 mm^3^, animals were randomized to treatment and control groups (10-11 mice / group) and treated p.o. with vehicle (40% Solutol HS-15, 10% ethanol, 50% water) or the in vivo probe BAY-593. Mice with HT-1080 tumors were treated with BAY-593 (2.5, 5, 10 or 20 mg/kg) with multiple dosing schedules (twice daily, BID; once daily, QD; every two days, Q2D; every four days, Q4D) for 14 days, whereas mice with MDA-MB-231 tumors were treated with BAY-593 (5 mg/kg, QD; 5 mg/kg, BID; 10 mg/kg, QD) for 12 days. Furthermore, the mice with subcutaneous PXF 541 tumors were treated with BAY-593 (5 mg/kg, QD; 10 mg/kg, QD) for 63 days. When PXF 541 tumors were approximately 90-100 mm^3^animals were randomized to test groups. Tumor response was assessed by measuring tumor volume [(length × width^2^/2)] using a caliper. Tumor volume and body weight were recorded 2-3 times weekly. Changes in body weight throughout the study compared to initial body weight at treatment start were considered as a measure of treatment-related toxicity (>20% weight loss = toxic). Tumor growth inhibition is presented as T/C ratio (treatment/control), calculated using tumor volumes at the end of the study.

### *ln vivo* compound exposure in the HT-1080 xenograft model

To quantify the concentrations of the drug candidate BAY-593 in circulation, BAY-593 was administered p.o. (per os, orally; n = 2-3 mice for each timepoint) in the respective efficacy study in a solubilized form. Plasma samples were taken from sacrificed animals at multiple timepoints after the last substance administration (e.g. 1, 3, 6, 24 hours). Samples were precipitated 1:5 (v:v) in ice-cold acetonitrile. After thawing and mixing, samples were centrifuged at 2000 × g at +4°C for 20 minutes and supernatants were analyzed by LC-MS/MS (AB Sciex). The concentrations of BAY-593 were determined using a calibration curve containing the same matrix as the injected sample and corrected for plasma protein binding. According to the free drug hypothesis (Smith et al., 2010), the unbound compound exposure in plasma is considered equal to the unbound concentration in the target tissue. The plasma concentration vs time (ct)-plots were determined for *in vivo* exposure. Area under the curve (AUC) was calculated according to the trapezoidal rule. HT-equilibrium dialysis was used to determine plasma protein binding using BAY-593 (3 µM) ^28^ and the unbound fraction (fu) was calculated.

### Statistical analyses

The preferred method for data analysis was ANOVA when the model assumptions were met. Methods used for assessing model fit included Shapiro-Wilk test for normality and Bartlett’s test for homogeneity of variances. Alternative analyses were a linear model with different variance terms for each group and finally non-parametric comparisons if the fit of both the parametric models was found lacking. Log-transformation was applied in cases where the distributions of measurements were heavily skewed. Nonparametric Mann-Whitney U-test was used for analyzing quantified TEAD-luc bioluminescence intensity. Statistical analyses were performed using the statistical programming language R (version 3.6.3). The validity of the model assumptions was checked. Group comparisons were corrected for false positive rate using either Tukey’s, Dunnett’s or Sidak’s method where appropriate. For YAP1/TAZ reporter assay and biochemical geranylgeranyl assay, IC_50_ values were calculated by 4-parameter fitting using a commercial software package (Genedata Screener). For YAP1/TAZ translocation assay, IC_50_ values were analyzed by four-parametric hill equation by Assay Analyzer software (Genedata).

## RESULTS

### Novel YAP1/TAZ pathway inhibitors identified via high-throughput screening

A high-throughput screen (HTS) using a library of 3.8 million small molecules was conducted to identify novel compounds that inhibit YAP1/TAZ activity. A reporter assay was performed using MDA-MB-231 human breast cancer cells expressing a YAP1/TAZ-dependent Firefly-luciferase reporter under control of TEAD-binding sites (TEAD-luciferase) and a constitutively active TK-Renilla-luciferase reporter as off-target control (**Figure 1A**). MDA-MB-231 cells contain a loss-of-function mutation in NF2 ^26^ an upstream component of the Hippo signaling pathway, leading to aberrant activation of YAP1/TAZ through constitutive nuclear localization and activation of YAP1/TAZ target genes. HTS hits that selectively inhibited TEAD-luciferase, but not TK-Renilla-luciferase were subsequently assessed regarding their ability to induce translocation of YAP1 from the nucleus to the cytoplasm, which is the physiological mechanism of inactivation. Finally, selected hits active in both assays were assessed regarding their effect on endogenous YAP1/TAZ target genes. These efforts led to the identification of BAY-856, an enantiomerically pure spiro-lactam stereoisomer (**Figure 1B**). BAY-856 demonstrated dose-dependent inhibition of TEAD-luciferase signal with an IC_50_ of 56 nM and no effect on TK-Renilla luciferase signal (**Figure 1C**). Moreover, BAY-856 effectively promoted YAP1 inactivation by translocation to the cytoplasm with an IC50 of 15 nM, assessed by high-content imaging. (**Figure 1D-E**). To confirm that the inhibitory activity of BAY-856 also translates to endogenous YAP1/TAZ target genes, we performed qPCR analysis on MDA-MB-231 cells treated with BAY-856. The well-established YAP1/TAZ target genes *ANKRD1*, *CTGF* and *CYR61* were down-regulated dose-dependently with IC50s of 81 nM, 76 nM and 86 nM, respectively, while no inhibition of control gene *ACTB* (IC50s > 10 µM) was observed. (**Figure 1F**). Overall, BAY- 856 demonstrated selective YAP1/TAZ inhibitory activity, which warranted further exploration of the unknown direct drug target.

**Figure 1.**
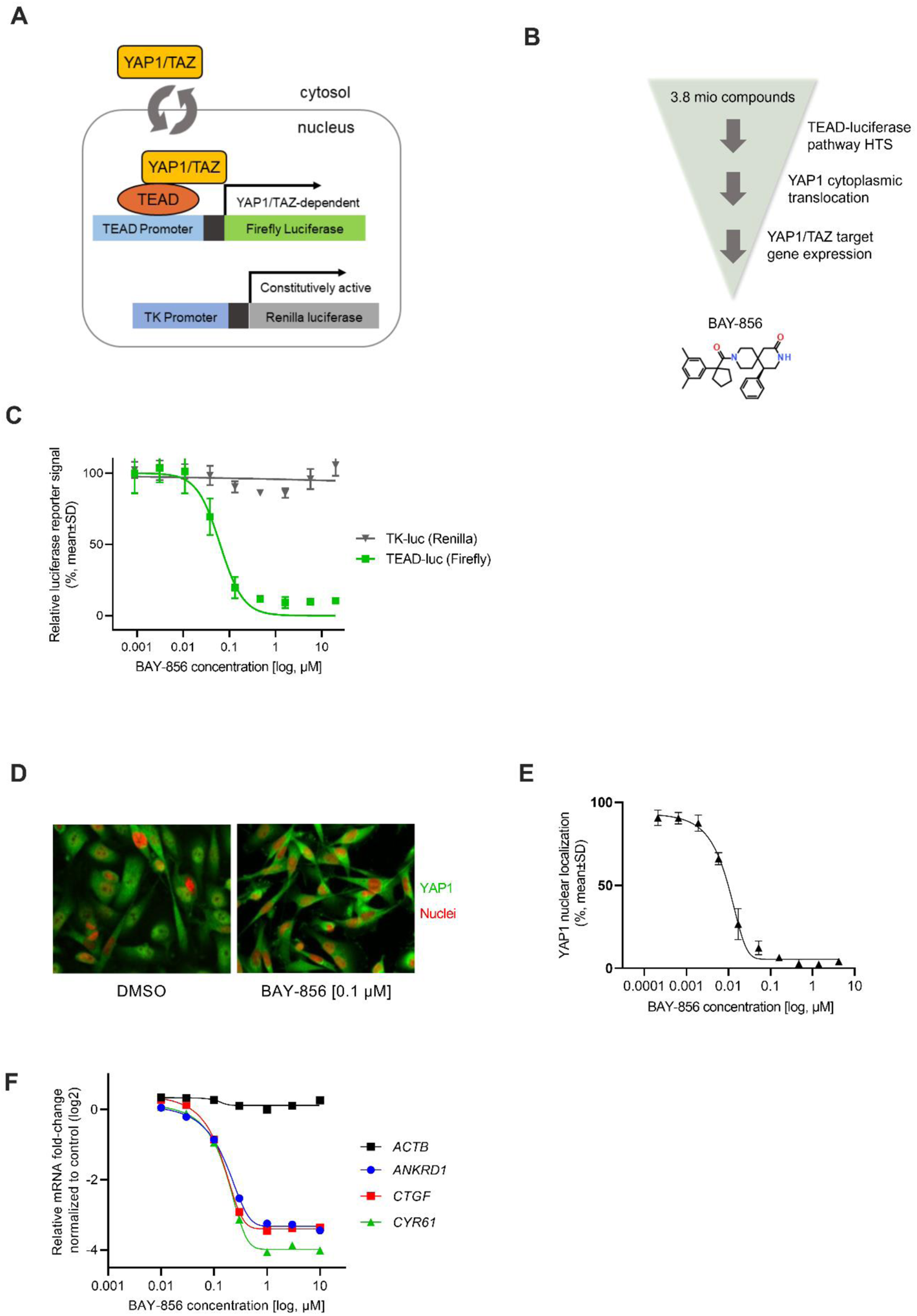
Novel YAP1/TAZ pathway inhibitors identified via high-throughput screen. (A) Schematic illustration of the YAP1/TAZ-luc dual reporter consisting of a TEAD promoter-YAP1/TAZ-dependent Luc2P firefly luciferase element and a constitutively active thymine kinase (TK) promoter-Renilla luciferase element expressed in MDA-MB-231 breast cancer cells. (B) Hit throughput screening cascade and structure of BAY-856 (C) BAY-856-induced inhibition of TEAD promoter-YAP1/TAZ-dependent Luc2P firefly luciferase (TEAD-luc FF) signal measured using the YAP1/TAZ reporter assay. The luciferase signal was normalized to vehicle-treated controls (100% activity) and thymidine kinase promoter-Renilla luciferase (TK-luc Ren) was used for toxicity control. Data are expressed as mean ± SD from three independent experiments performed in duplicate. (D) YAP1/TAZ translocation from the nucleus to the cytoplasm in MDA-MB-231 cells treated with BAY-856 [0.1 µM] and DMSO control. YAP1 (green) and nucleic acid (red) staining was determined using high-content analysis (HCA) microscopy. (E) Dose response curve analysis of YAP1/TAZ translocation from the nucleus to the cytoplasm in MDA-MB-231 cells treated with BAY-856 and DMSO control. YAP1 (green) and nucleic acid (red) staining was determined using high-content analysis (HCA) microscopy. Data are expressed as mean ± SD from three independent experiments performed in duplicate. (F) Dose response fold change values of YAP1/TAZ target genes ANKRD1, CTGF, CYR61 and the housekeeping genes *ACTB*, *GAPDH* in BAY-856-treated (0.01-10 μM) MDA-MB-231 cells compared with vehicle-treated control cells. Relative mRNA expression was determined by quantitative PCR.

### PGGT1B, a subunit of the GGTase-I complex, is the direct target of the novel YAP1/TAZ pathway inhibitors

To identify the direct target of BAY-856, proteome-wide cellular thermal shift assays combined with mass spectrometry (CETSA-MS) were used. CETSA-MS allows for the identification of direct binding partners of small molecules ^29^. These studies identified geranylgeranyltransferase type-1 subunit β (PGGT1B) as the top-ranked hit for direct binding of BAY-856, inducing a shift in the melting temperature of PGGT1B between the control treatment and BAY-856 (**Figure 2A**). A similar result was observed for BAY-6092, a close analog of BAY-856 with improved cellular potency. (**Figure S1A)**. Direct PGGT1B binding by BAY-856 was confirmed by thermal shift assay (TSA) using recombinant PGGT1B, leading to a shift of 12.9 K in melting temperature (ΔTM) upon treatment compared to DMSO-treated control (**Figure 2B**). To assess potential off-target kinase inhibitory activity of BAY-856, a panel consisting of 337 kinases was assessed (Eurofins Discovery). BAY-856 did not display any relevant activity when tested a 1 µM (**Supplementary Table S1**).

**Figure 2.**
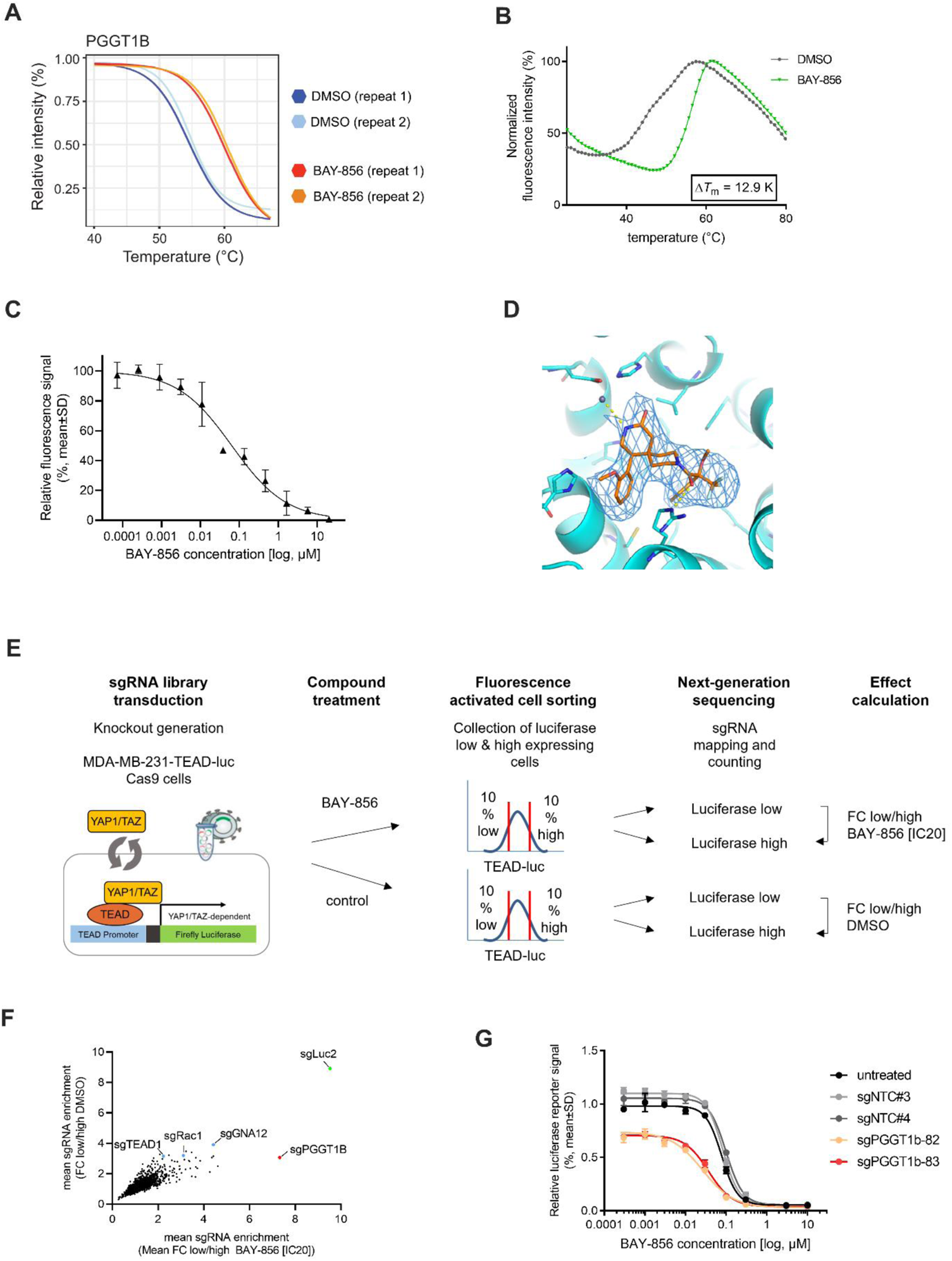
PGGT1B, a subunit of the GGTase I complex, is the direct target of the novel YAP1/TAZ pathway inhibitors. (A) BAY-856 target engagement in MDA-MB-231 cell lysates treated with 10 μM BAY-856 or vehicle control as determined by proteome-wide cellular thermal shift assay mass spectrometry (CETSA MS). Fold change values were normalized to a constant value of 1 by using the lowest temperature condition (+40°C) as reference. (B) Direct PGGT1B binding by BAY-856 by thermal shift assay (TSA) using recombinant PGGT1B. (C) BAY-856-induced inhibition of human GGTase-I activity measured using the biochemical GGTase-I assay. Fluorescence signal intensity is normalized to both DMSO control (0% signal intensity) and the control sample (100% signal intensity) containing complete reaction without either enzyme or BAY-856. Data are expressed as mean ± SD. (D) X-ray co-crystallization of the complex formed by the purified rat GGTase protein and the GTTase-I inhibitor BAY-6092. (E) Schematic presentation of strategy for pooled CRISPR/Cas9 YAP1/TAZ pathway screen to aid target deconvolution of BAY-856. (F) Using a library of sgRNAs targeting approximately 6000 human genes, MDA-MB-231 cells were treated with BAY-856 (IC_20_) or left untreated. Data is presented as ratios of mean fold change of sgRNAs targeting genes (BAY-856 vs untreated). The cells were sorted as “luciferase high” or “luciferase low” based on their luciferase expression. sgLuc2 represents a custom made sgRNA against the TEAD-luc reporter gene, which was spiked into the sgRNA library as positive control. (G) IC_50_ values for BAY-856 for individual sgRNAs against *PGGT1B* (sgPGGT1B-82 and sgPGGT1B-83), non-targeting sgRNAs (sgNTC#3, sgNTC#4) and untreated control were determined by measuring luciferase activity (relative luciferase unit, RLU). The figure demonstrates a response curve of MDA-MB-231 TEAD-luc Cas9 cells treated for 24 hours with increasing concentrations of BAY-856. Data are expressed as mean ± SD.

PGGT1B is a subunit of geranylgeranyltransferase type-1 (GGTase-I), an enzyme required for the addition of a 20-carbon geranylgeranyl moiety to proteins containing a CaaX motif. Therefore, the effect of BAY-856 on GGTase-I was investigated using a suitable biochemical assay. Based on the activity of GGTase-I to transfer a geranylgeranyl moiety from geranylgeranyl pyrophosphate (GGPP) to a peptide containing the recognition sequence GCVLL resulting in fluorescent signal, GGTase-I activity was shown to be inhibited dose-dependently by BAY-856 (**Figure 2C**). To gain further insights into the binding mode, X-ray crystallography was used to determine the crystal structure of GGTase-I in complex with BAY-6092. The co-crystal structure was determined at 3.67 Å using X-ray crystallography with a crystal form containing six protein dimer molecules per asymmetric unit (**Figure 2D**). In summary, PGGT1B, a subunit of the GGTase-I complex, was validated to be the direct target of the novel YAP1/TAZ pathway inhibitors.

To independently confirm PGGT1B as the target of BAY-856, a CRISPR/Cas9 knockout (KO) screen was performed in MDA-MB-231 TEAD-luc Cas9 cells. (**Figure 2E**). The cells were stained with a specific anti-firefly luciferase antibody suitable to detect dose-dependent differences in TEAD-luciferase activity upon BAY-856 treatment using FACS (**Figure S1B**). Using a high coverage pooled library of sgRNAs (average 8 sgRNAs/target) targeting approximately 6000 human genes and 200 negative and positive control sgRNAs, several sgRNAs targeting *PGGT1B,* amongst others, were found to be markedly enriched in the luciferase low population of the untreated cell pool, indicating that attenuating endogenous *PGGT1B* levels via CRISPR/Cas9 inhibits cellular TEAD-luciferase activity (**Figure 2F**). Notably, low dose (IC_20_) BAY-856 treatment further enhanced the enrichment in the luciferase low population only in cells targeted by *PGGT1B* sgRNAs (**Figure 2F**), indicating that attenuating endogenous *PGGT1B* levels in CRISPR-sgRNA targeted cells sensitized these cells to BAY-856.

To further validate these results, individual sgRNAs against *PGGT1B* were investigated for their ability to increase sensitivity to BAY-856. Two individual sgRNAs (sgPGGT1B-82 and sgPGGT1B-83) led to marked reduction of PGGT1B protein levels in a pool of transduced cells, decreasing the luciferase activity in the presence of BAY-856 compared to non-targeting sgRNAs (sgNTC#3, sgNTC#4) or untreated control (**Figure 2G**). These results were validated by immunoblotting, showing knockout of PGGT1B protein in *PGGT1B*-targeted cell pools transduced with the same specific *PGGT1B* targeting sgRNAs (**Figure S1C**). These CRISPR screening results strongly supported the finding that *PGGT1B* is the molecular target of BAY-856.

### BAY-856 blocks YAP1/TAZ activity via blockade of Rho-GTPase signaling

Having identified GGTase-I as the direct target, we reasoned that Rho-GTPase activity might be affected by BAY-856. The activity of Rho-GTPases relies on their ability to localize to the plasma membrane, a process dependent on transfer of a geranylgeranyl moiety to a carboxy-terminal cysteine residue by the GGTase-I. Moreover, Rho-GTPases are well-known positive upstream inputs for YAP1/TAZ activity by modulation of the actin cytoskeleton ^30,31^.

To assess the amount of membrane-bound, active Rho-GTPases, we used cell fractionations of BAY-856 treated cells followed by protein capillary electrophoresis. The amount of RhoA, RhoC, Rac1, Rap1A/B and Cdc42 at the membrane of MDA-MB-231 cells was markedly reduced by BAY-856 treatment, while proteins of Ras-GTPase family (RalA, RalB and Ras) remained unaffected by BAY-856 treatment (**Figure 3A**).

**Figure 3.**
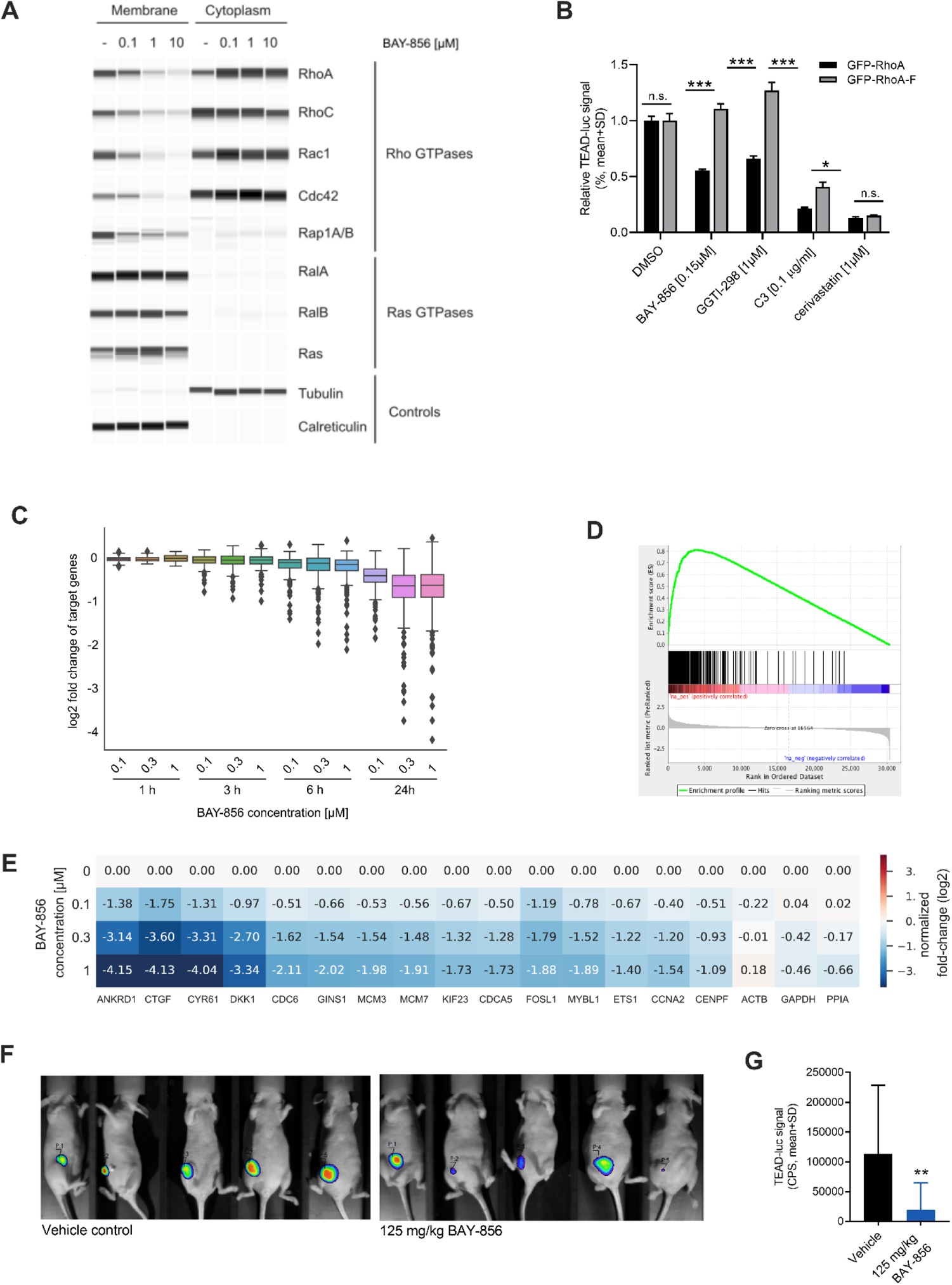
BAY-856 inhibits YAP1/TAZ signaling geranylgeranyl-dependently. (A) The effect of BAY-856 [0.1, 1 or 10 μM] on Rho- and Ras-GTPase protein localization in MDA-MB-231 cells measured by capillary electrophoresis. Calreticulin and tubulin were used as controls for GTPase proteins measured at the plasma membrane and in cytoplasmic fractions, respectively. (B) Relative TEAD-luc signal in GFP-RhoA- or GFP-RhoA-F-expressing MDA-MB-231 cells treated with 150 nM BAY-856, 2 nM BAY-6092, 1 μM GGTI-298, 0.1 μg/mL C3, 1 μM cerivastatin or vehicle for 24 hours, as determined using the geranylgeranyl reporter assay. Data are expressed as mean + SD. ANOVA: *, p<0.05; ***, p<0.001. (C) mRNA expression of 379 YAP1/TAZ target genes in MDA-MB-231 cells treated with BAY-856 (0.1, 0.3 or 1 µM) or vehicle control for 1, 3, 6 or 24 hours, determined by RNA-sequencing. (D) Gene set enrichment analysis of a YAP1/TAZ gene signature consisting of 379 YAP1/TAZ target genes in MDA-MB-231 cells treated with 0.3 µM BAY-856 for 24 hours. (E) Fold change values (log2) of selected YAP1/TAZ target genes and the housekeeping genes *ACTB*, *GAPDH* and *PPIA* in BAY-856-treated (0.1, 0.3 or 1 µM) MDA-MB-231 cells compared with vehicle-treated control cells. Relative mRNA expression was determined by quantitative PCR. (F) Representative *in vivo* bioluminescence images of MDA-MB-231 TEAD-luc tumor-bearing mice on treatment day 2. The mice were treated with vehicle (n=8) or 125 mg/kg BAY-856 (n=9) twice daily for 10 days. Scalebars indicate TEAD-luc signal intensity. (G) Quantified TEAD-luc bioluminescence signals determined from the MDA-MB-231 TEAD-luc tumors described in (F). Data are expressed as mean + SD. CPS, counts per second. Mann-Whitney U-test: **, p=0.005.

To show that inhibition of YAP1/TAZ activity via Rho-GTPases is geranylgeranylation-dependent, the effect of BAY-856 on TEAD-luciferase activity were assessed in MDA-MB-231 cells expressing either GFP-RhoA containing a natural geranylgeranylation motif or a modified GFP-RhoA-F bearing a consensus site for farnesylation. Independent of geranylgeranylation inhibition, GFP–RhoA-F localizes on cell membrane and sustains YAP1/TAZ nuclear localization and transcriptional activity (Sorrentino et al., 2014), thus serving as a specificity control for GGTase-I inhibitors. BAY-856 at 0.15µM was shown to induce inhibition of the TEAD-luc reporter comparable to inhibition by 1 μM of the GGTase-I inhibitor GGTI-298. Using these concentrations, the TEAD-luc reporter signal was reduced by BAY-856 and GGTI-298 in MDA-MB-231 cells transfected to express GFP-RhoA, whereas in cells transfected to express GFP-RhoA-F, the farnesylation motif was sufficient to rescue YAP1/TAZ activation and maintain transcriptional activity in the presence of these compounds (**Figure 3B**). As expected, C3 and cerivastatin inhibited the TEAD-luc reporter irrespectively of GFP-RhoA or GFP-RhoA-F. The protein expression of GFP-RhoA or GFP-RhoA-F in the transfected MDA-MB-231 cells was confirmed by protein immunoblotting (**Figure S2A**).

Having established the connection between Rho GTPase inhibition and YAP1/TAZ activity, we next applied global gene expression profiling by RNA sequencing (RNA-seq). We found that the set of 379 previously described YAP1/TAZ target genes in MDA-MB-231 cells ^25^ was significantly downregulated in a dose- and time-dependent manner by BAY-856 treatment (**Figure 3C, Figure S2B**). This observation was further supported by a gene set enrichment analysis, which identified the YAP1/TAZ gene signature as the most significantly differentially expressed gene set. (**Figure 3D**). To validate the RNA-seq results, mRNA expression of YAP1/TAZ target genes was analyzed in MDA-MB-231 cells using quantitative PCR. In line with the RNA-Seq results, the known YAP1/TAZ target genes *ANKRD1, CYR61, CTGF, DKK1, CDC6, GINS1, MCM3, MCM7, KIF23, CDCA5, FOSL1, MYBL1, ETS1, CCNA2* and *CENPF* were downregulated by BAY-856 treatment. (**Figure 3E**).

To demonstrate the inhibitory effect of BAY-856 on YAP1/TAZ signaling *in vivo*, MDA-MB-231 TEAD-luc cells were inoculated subcutaneously into the flank of nude mice. The mice were treated daily with 125 mg/kg BAY-856 or vehicle for 3 days and subjected to *in vivo* bioluminescence imaging (**Figure 3F**). The bioluminescence signal intensity was reduced in mice treated with BAY-856 (p=0.005) compared to vehicle-treated mice, demonstrating that BAY-856 treatment resulted in decreased intra-tumoral YAP1/TAZ signaling (**Figure 3G**).

### Pharmacological inhibition of YAP1/TAZ signaling impairs cancer cell proliferation in vitro

YAP1/TAZ regulate cancer cell proliferation and invasion through activation of downstream targets involved in cell cycle progression. To assess the effect on cancer cell proliferation, we used BAY-6092, a close analog of BAY-856 with improved cellular potency **(Figure 4A**, **Table 1**). We independently confirmed that the anti-proliferative activities of BAY-6092 and BAY-856 were conserved. A comparison of the IC50 values for both compounds highlighted the increased potency of BAY-6092 and resulted in a good correlation among cell lines tested (Pearson r = 0.95) **(Figure 4B**). BAY-6092 inhibited cell proliferation defined by an activity area >4 in 37 out of 250 cell lines, with the strongest activity observed in cell lines derived from soft tissue sarcomas, thyroid, breast, kidney, pancreas, prostate and head and neck cancers **(Figure 4C**). Activity areas and IC_50_ values for each cell line are summarized in **Supplementary Table 3**.

**Figure 4.**
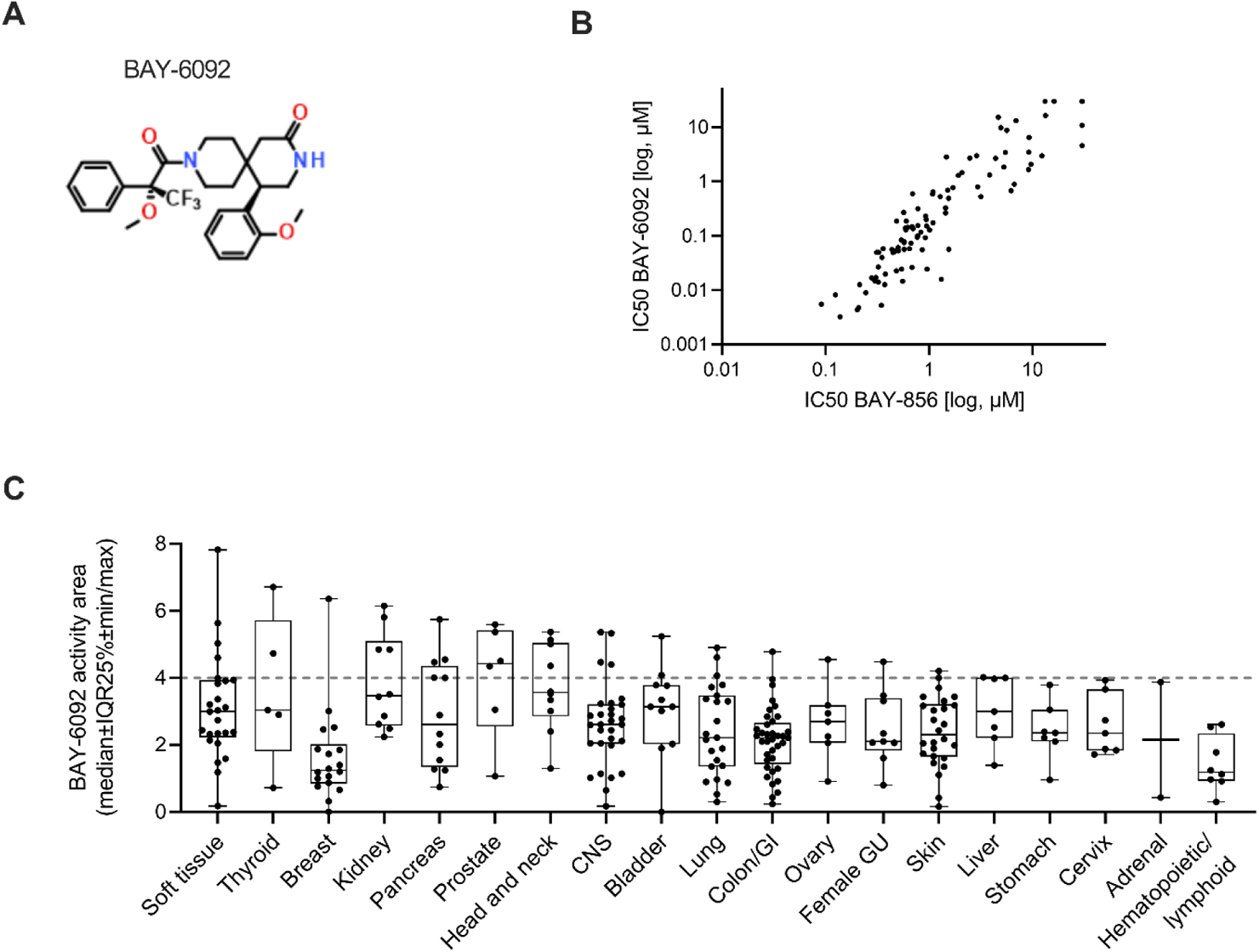
Pharmacological inhibition of YAP1/TAZ signaling impairs cell proliferation *in vitro*. (A) Structure of BAY-6092 (B) Correlation of anti-proliferative activity (IC50 values) of BAY-586 and BAY-6092 based on cancer cell line screening. (C) Cell proliferation as activity area in the OncoPanel™ Cell Proliferation assay screen of 250 human cancer cell lines treated with BAY-6092 for 72 hours. The cell lines were categorized based on the activity area of each cancer type and ranked accordingly, each dot presenting the activity area of a single cell line. Box plots describe median, 25/75% quartiles and minimum and maximum values. Cell lines above grey dashed line (activity area >4) are considered sensitive to BAY-6092 treatment.

**Table 1:**
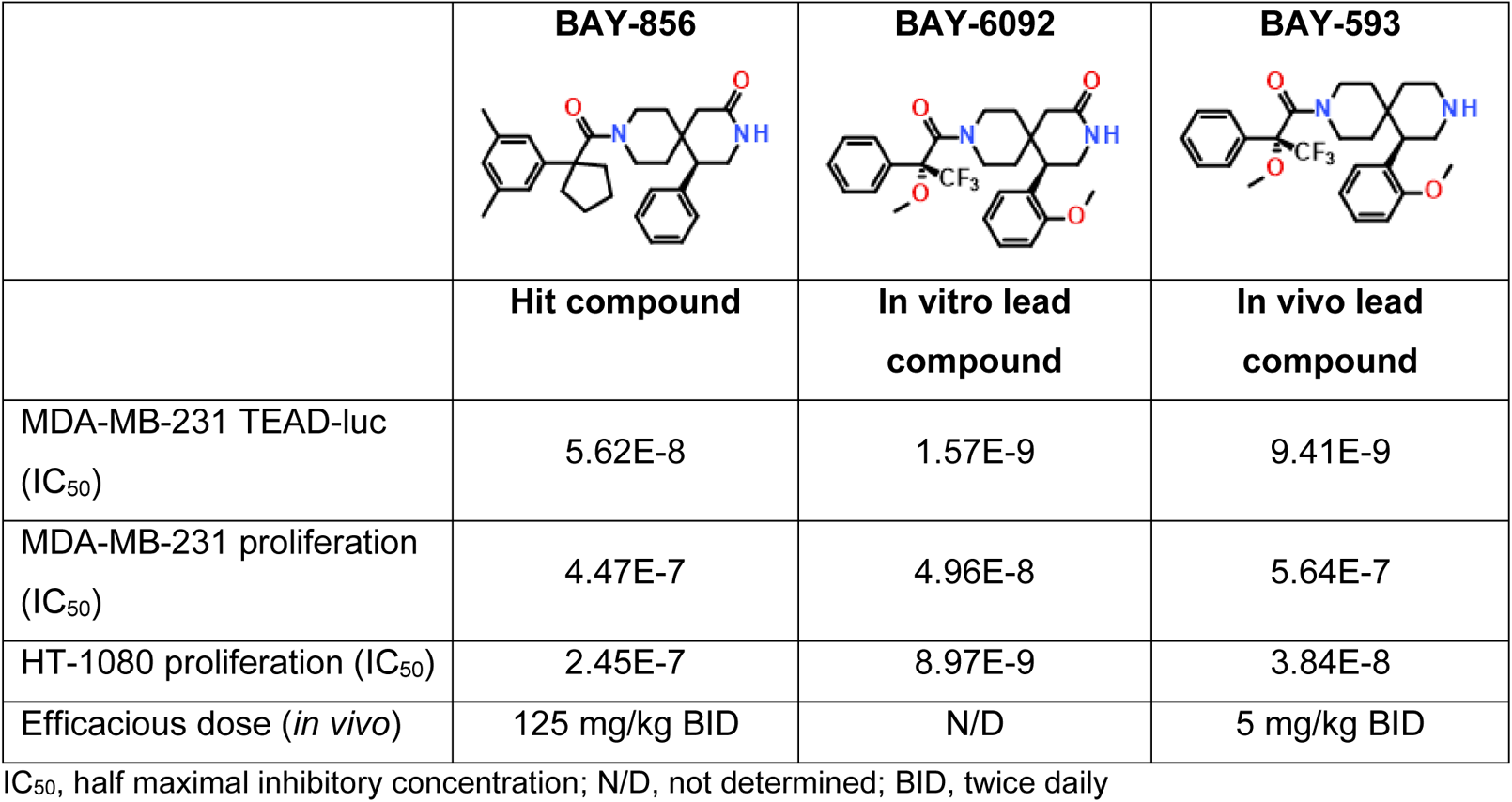
In vitro and in vivo properties of novel small molecule inhibitors.

|  | **BAY-856** | **BAY-6092** | **BAY-593** |
| --- | --- | --- | --- |
|  | **Hit compound** | **In vitro lead compound** | **In vivo lead compound** |
| MDA-MB-231 TEAD-luc (IC<sub>50</sub>) | 5.62E-8 | 1.57E-9 | 9.41E-9 |
| MDA-MB-231 proliferation (IC<sub>50</sub>) | 4.47E-7 | 4.96E-8 | 5.64E-7 |
| HT-1080 proliferation (IC<sub>50</sub>) | 2.45E-7 | 8.97E-9 | 3.84E-8 |
| Efficacious dose (*in vivo*) | 125 mg/kg BID | N/D | 5 mg/kg BID |

IC<sub>50</sub>, half maximal inhibitory concentration; N/D, not determined; BID, twice daily

### Pharmacological inhibition of YAP1/TAZ signaling inhibits tumor growth *in vivo*

To assess the therapeutic benefit of YAP1/TAZ signaling blockade *in vivo*, we chose the *in vivo* lead compound BAY-593 (**Figure 5A**), which was developed from systematic optimization studies focusing on improving the *in vitro* and *in vivo* pharmacokinetic profile of BAY-6092 (**Table 1**). To ensure translatability of the results obtained with the *in vitro* lead compound BAY-6092, we confirmed that the anti-proliferative activities of BAY-6092 and BAY-593 were conserved. A comparison of the IC50 values for both compounds resulted in a good correlation among cell lines tested (Pearson r = 0.97) (**Figure S4F**) The *in vivo* anti-tumor activity of BAY-593 was evaluated in the HT-1080 fibrosarcoma and MDA-MB-231 triple-negative breast cancer (TNBC) xenograft mouse models, as these were identified as responders in the cell line screen. BAY-593 was dosed at 2.5-20 mg/kg, p.o. using daily and intermittent dosing regimens: twice daily (BID), once daily (QD), every two days (Q2D) and every four days (Q4D). In the HT-1080 model, treatment with BAY-593 resulted in dose-dependent anti-tumor activity (**Figure 5B**). In the MDA-MB-231 xenograft model, BAY-593 also demonstrated anti-tumor activity at all dosing schedules tested (5 mg/kg, BID or QD; 10 mg/kg QD) (**Figure 5C**). BAY-593 treatment resulted in a strong dose-dependent reduction of the HT-1080 tumor volumes on day 9 (**Figure 5D, Figure S3A**) and also reduced the MDA-MB-231 tumor volumes on day 12 (**Figure 5E**, **Figure S3B**) in comparison to vehicle-treated tumors. In addition, total eradication of HT-1080 tumor was observed in 4 out 10 mice when BAY-593 was dosed at 10 mg/kg QD (**Figure 5D**). All doses and schedules tested were well-tolerated in the HT-1080 and MDA-MB-231 models with a maximum body weight loss of less than 10% (**Figure S3C-D**). To confirm the downregulation of YAP1/TAZ target genes identified by RNA-Seq of BAY-856-treated cells *in vitro*, HT-1080 tumor bearing-mice were treated with a single dose of BAY-593 (2.5-20 mg/kg) and tumors were collected 6 hours after dosing. The collected tumors were analyzed by qRT-PCR. A dose-dependent downregulation in the expression of YAP1/TAZ target genes (*CYR61, CTGF, ANKRD1, CD274*) was observed in tumors treated with BAY-593 compared to vehicle treated animals (**Figure 5F**).

**Figure 5.**
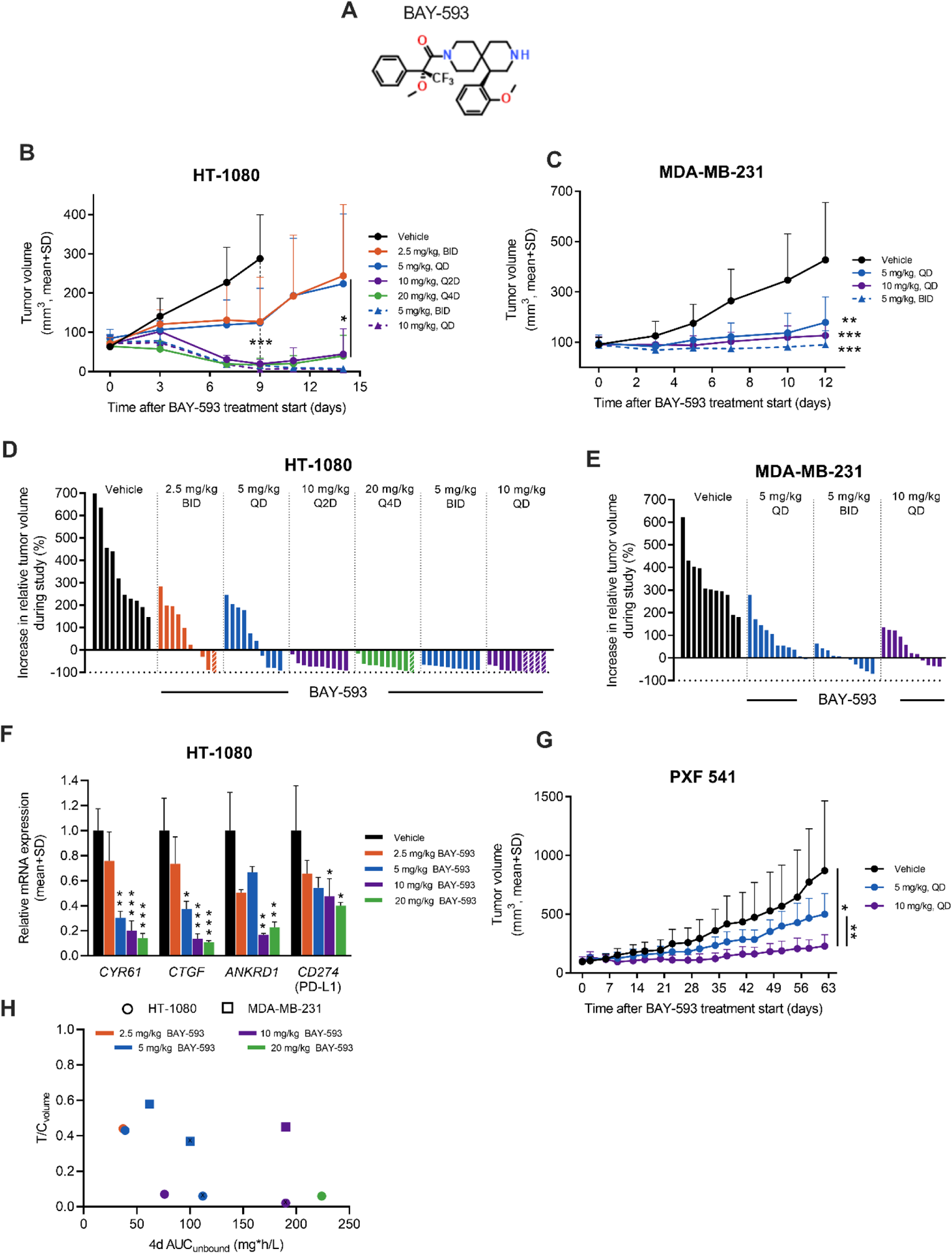
YAP1/TAZ pathway inhibitor BAY-593 reduces tumor growth *in vivo*. (A) Structure of BAY-593 (B) Growth curves and (C) increase in relative tumor volumes of HT-1080 human fibrosarcoma tumors in female NMRI nu/nu mice (n=10/group) treated with BAY-593 (2.5, 5, 10 or 20 mg/kg, p.o) at the indicated dosing schedules (twice daily, BID; once daily, QD; every two days, Q2D; every four days, Q4D) for 14 days. Vehicle-treated mice were treated for 9 days (black line in the panel A). Striped columns indicate mice with total eradication of tumor (increase in relative tumor volume = −100%) in the panel C. Variance was estimated separately for each group using weighted least squares: *, p<0.05; ***, p<0.001. (D) Growth curves and (E) increase in relative tumor volumes of MDA-MB-231 human breast cancer tumors in female NMRI nu/nu mice (n=11/group) treated with vehicle or BAY-593 (5 mg/kg, QD, p.o.; 5 mg/kg, BID, p.o.; 10 mg/kg, QD, p.o.) for 12 days. ANOVA in log scale: **, p<0.01; ***, p<0.001. (F) Relative mRNA expression levels of the YAP1/TAZ target genes *CYR61, CTCF, ANKRD1* and *CD274* (coding PD-L1) in HT-1080 tumors (n=3/group) collected from mice treated with vehicle or a single dose of BAY-593 (2.5, 5, 10 or 20 mg/kg, p.o) 6 hours before sacrifice. Relative mRNA levels were normalized to the corresponding gene expression levels in vehicle-treated tumors. For statistics, ANOVA and variance estimated separately for each group using weighted least squares were used for *CYR61*/*CD274* and *CTCF/ANKRD1*, respectively. *, p<0.05; **, p<0.01, ***, p<0.001, compared to vehicle. (G) Growth curves of PXF 541 pleuromesothelioma PDX tumors in female NMRI nu/nu mice (n=10/group) treated with vehicle or BAY-593 (5 mg/kg, QD, p.o.; 10 mg/kg, QD, p.o.) for 63 days. Variance was estimated separately for each group using weighted least squares: *, p=0.01; **, p=0.003. (H) *In vivo* exposure of BAY-593 expressed as AUC levels during the 4-day cycle (4d AUC) in relation to the Treatment/Control (T/C_volume_) ratios determined from HT-1080 and MDA-MB-231 tumors on day 9 or day 12, respectively. Dots and squares demonstrate HT-1080 or MDA-MB-231 tumor-bearing mice, respectively.

To test whether BAY-593 inhibits tumor growth in a patient-derived xenograft (PDX) model, the PXF 541 pleuromesothelioma model was selected due to its previously identified sensitivity in an *ex vivo* 3D tumor colony assay (data not shown). In the PXF 541 model, BAY-593 treatment with 5 mg/kg QD and 10 mg/kg QD resulted in dose-dependent anti-tumor activity *in vivo* (**Figure 5G**). Both schedules were well-tolerated with acceptable weight loss (approximately 10%) (**Figure S3E**).

To gain insights into the pharmacokinetic-pharmacodynamic (PK-PD) relationship, the in vivo exposure of BAY-593 in the HT-1080 and MDA-MB-231 models was investigated. The observed anti-tumor efficacies showed a dependency on the exposures reached in mice after treatment with different dosing regimens (**Table 2**). Particularly in the HT-1080 model, high AUC (area under the curve) values during the 4-day cycle (4d AUC) were associated with a more pronounced decrease in T/C_max_ ratio on day 15, independently of the dosing schedule (**Figure 5H**). AUC was therefore identified as the driver of efficacy for BAY-593.

**Table 2:** *In vivo* pharmacokinetics of BAY-593 in the HT-1080 and MDA-MB-231 xenograft models.

| **Dose** | **Schedule** | **T/C<sub>volume</sub>** | **C<sub>max</sub>** | **4d AUC** | **4d AUC<sub>unbound</sub>** |
| --- | --- | --- | --- | --- | --- |
| (mg/kg) |  | on day 9 | (µg/L) | (mg*h/L) | (µg*h/L) |
| 2.5 | BID | 0.44 | 100 | 2.6 | 37 |
| 5 | QD | 0.43 | 76 | 2.8 | 39 |
| 10 | Q2D | 0.069 | 350 | 5.4 | 76 |
| 20 | Q4D | 0.063 | 2262 | 16.0 | 224 |
| 5 | BID | 0.056 | 275 | 8.0 | 112 |
| 10 | QD | 0.021 | 321 | 14.0 | 196 |

**MDA-MB-231**
| **Dose** | **Schedule** | **T/C<sub>volume</sub>** | **C<sub>max</sub>** | **4d AUC** | **4d AUC<sub>unbound</sub>** |
| --- | --- | --- | --- | --- | --- |
| (mg/kg) |  | on day 12 | (µg/L) | (mg*h/L) | (mg*h/L) |
| 5 | QD | 0.58 | 176 | 4.4 | 62 |
| 5 | BID | 0.37 | 313 | 7.1 | 100 |
| 10 | QD | 0.45 | 406 | 14 | 190 |

4d AUC, area under the curve during the 4-day cycle; BID twice daily; C<sub>max</sub>, maximal (peak) plasma concentration; QD, once daily; Q2D, every two days; Q4D, every four days; T/C<sub>volume</sub>, treatment/control ratio of tumor volume.

## DISCUSSION

The results presented in the study highlight the identification and characterization of novel inhibitors targeting the YAP1/TAZ pathway. We show that phenotypic screening is a viable strategy to identify novel small molecules when combined with a matching target deconvolution strategy. We identified several hit classes in the HTS and used a screening tree of secondary YAP1/TAZ assays to select the most promising hit cluster for target deconvolution and lead optimization. BAY-856 was identified as the hit with selective activity in all assays.

To identify the direct target of BAY-856, the target deconvolution strategy consisted of complementary biochemical, biophysical and functional genomics methods, leading to the identification of GGTase-I as the direct target of BAY-856. We show that CETSA-MS is a powerful tool to identify small molecule binders when combined with classical thermal shift (TSA) and biochemical assays ^32^. Finally, X-ray crystallography demonstrated that despite the moderate resolution, the omit electron density obtained for the ligand clearly revealed the binding mode of BAY-6092 and showed that its binding site overlapped to a significant extent with the geranylgeranyl-diphosphate recognition site.

It has previously been shown that functional genomics screening can be used to identify potential functional molecular targets of small molecules ^33^. Therefore, to support the target deconvolution for BAY-856, a systematic, large scale CRISPR/Cas9 screen in combination with focused small scale functional genomic screening of YAP1/TAZ regulators was used. The logic behind the screens was to identify factors that upon attenuation could sensitize cells towards BAY-856 inhibition. Strikingly, large-scale as well as focused small-scale functional genomic screening applying CRISPR/Cas9 or RNAi technology resulted in pronounced sensitization towards BAY-856 inhibition exclusively when PGGT1B expression was perturbated. While the homozygous knock out of PGGT1B most likely mimics a complete inhibition of PGGT1B by BAY-856 and will thus not contribute to the observed sensitization phenotype, the imperfect heterozygous knock out in some cells results in reduced expression of functional PGGT1B protein and can therefore explain the observed sensitization of PGGT1B expression perturbated cells in both CRISPR/Cas9 screens as well as in the pool of cells edited with individual sgRNAs against PGGT1B.

We further explored the mechanism of action of BAY-856 and its impact on YAP1/TAZ signaling. YAP1/TAZ are key regulators of mechanotransduction by translating extracellular matrix (ECM) cues to the nucleus ^34^. It has previously been shown that GGTase-I inhibitors block YAP1/TAZ activity via Rho-GTPase signaling ^27,30,31^. Rho-GTPase function depends on a specific post-translational modification, the transfer of a geranylgeranyl moiety to a carboxy-terminal cysteine residue, leading to Rho-GTPase component recruitment and lipid-anchoring to the plasma membrane ^35^. This dependency was demonstrated by BAY-856 treatment, resulting in the plasma membrane delocalization of Rho-GTPase components while the expression levels of them in cytoplasm remained unaffected as seen by immunoblotting. In the absence of activated/membrane-bound Rho-GTPases, for instance when geranylgeranylation of Rho-GTPases is blocked by GGTase-I inhibition ^36^, the transduction of activated mechanical stimuli to YAP1/TAZ is impaired, resulting in the inhibition of YAP1/TAZ transcriptional and biological responses ^30^. To verify whether the inhibition of GGTase-I is part of the molecular mechanism of action by BAY-856 and BAY-6092, the effects of these compounds were studied in MDA-MB-231 cells expressing either GFP-RhoA or GFP-RhoA-F, bearing a natural geranylgeranylation motif or a motif for farnesylation, respectively. GFP-RhoA-F has been shown to localize at the plasma membrane and sustain YAP1/TAZ activity also in the presence of the GGTase inhibitor GGTI-298 ^27^, confirming that farnesylation promotes membrane tethering of GFP-RhoA-F independently of geranygeranyltransferases. In a geranylgeranyl reporter assay, both BAY-856 and BAY-6092 inhibited geranylgeranyltransferases either directly or indirectly and impaired RhoA subcellular localization. However, control compounds, C3 ^26^ and cerivastatin ^27^, inhibited the YAP1/TAZ activity both in the presence of GFP-RhoA and GFP-RhoA-F, because C3 inhibits Rho-GTPases by ADP-ribosylation whereas cerivastatin inhibits the production of metabolites required for both geranylation and farnesylation.

Rho-GTPases regulate various cellular mechanisms, including cell migration, proliferation, and survival. To assess the contribution on YAP1/TAZ inhibition, we profiled global effects on gene expression by RNA sequencing of MDA-MB-231 breast cancer cells treated with BAY-856. We could show that the previously identified YAP1/TAZ target genes ^25^ are the most significantly downregulated set of genes, suggesting that YAP1/TAZ are mayor downstream effectors of Rho-GTPase signaling. Moreover, we demonstrate that BAY-856 can block YAP1/TAZ activity using the TEAD-luc reporter in MDA-MB-231 xenografts in vivo.

To assess the broader therapeutic potential of the novel YAP1/TAZ pathway inhibitors, we profiled the anti-proliferative activity of BAY-6092, an analog of BAY-856 with improved in vitro potency, in a panel of 250 cancer cell lines. Soft tissue sarcomas, thyroid, breast, kidney, pancreas, prostate and head and neck cancers were the indications with the strongest enrichment in responder cell lines. While breast cancer cell lines mostly did not respond to BAY-6092, NF2-null (Hippo pathway altered) MDA-MB-231 ^37^ cells were found to be sensitive to BAY-6092.

Several GGTase-I inhibitors have been developed over the years ^38,39^. The previously published GGTase-I inhibitors are characterized by challenging in vivo PK properties and the need for i.p. or i.v. infusion dosing ^40^. Currently, only one geranylgeranyltransferase-1 inhibitor known as GGTI-2418 (PTX-100) has advanced to early clinical trials (NCT03900442). GGTI-2418 was safe and tolerated at all doses tested with some evidence of disease stability. Insights into the effect of GGTase-I inhibition were nevertheless limited due to the short half-life of the drug ^41^. A Phase 1 pharmacodynamic and pharmacokinetic basket study involving cancer patients with advanced solid and hematological malignancies, including cases with relapsed/refractory PTCL or multiple myeloma (MM) is currently in progress ^42^. Our lead molecule BAY-593 has been optimized for favorable in vivo properties compatible with oral dosing. BAY-593 achieved potent anti-efficacy in cell-line-derived (CDX) as well as patient-derived xenograft models, including complete remissions in the HT-1080 xenograft model. Interestingly, the HT-1080 cell line contains an oncogenic mutation in Rac1 (N92I) ^43^, which may explain the sensitivity of this cell line towards BAY-593. Moreover, we were able to confirm the downregulation of YAP1/TAZ target genes previously identified by RNA-Seq in vivo, confirming blockade of the pathway in vivo. CYR61, CTGF, ANKRD1 therefore may serve as pharmacodynamic biomarker candidates of response to BAY-593. The down-regulation of CD274 (encoding PD-L1) is furthermore in line with previously published results showing YAP1 as a regulator of PD-L1 expression in cancer ^44^. We further sought to identify the most efficacious in vivo dosing schedule for BAY-593. To this end, mechanistic modelling of the PK/PD relationship was used to identify AUC as the driver of efficacy of BAY-593 in vivo.

In conclusion, we show here the identification and pharmacological characterization of novel YAP1/TAZ pathway inhibitors with potent anti-tumor activity via blockade of GGTase-I / Rho-GTPase signaling. Our lead molecule BAY-593 is a novel tool compound to explore Rho-GTPase signaling and downstream YAP1/TAZ biology in vitro and in vivo. Further investigations and preclinical studies are warranted to explore the full therapeutic potential of these inhibitors and their efficacy in different cancer types.

The therapeutic concept of small molecule modulation of protein lipidation demonstrated here has successfully been shown for other targets. For example, small molecule inhibitors of protein farnesylation have been developed for many years and recently, the small molecule Lonafarnib has been approved for the treatment of Hutchinson-Gilford Progeria Syndrome^45^. In fact, an alternative strategy for inhibition of YAP1/TAZ activity via lipid modulation may be achieved via blockade of TEAD protein auto-palmitoylation. In preclinical models, these small molecules have been shown to disrupt YAP1/TAZ-TEAD interaction and induce anti-tumor effects in vivo ^46^. Moreover, in a recent phase 1 study (NCT04665206), the TEAD auto-palmitoylation Inhibitor VT3989 was shown to be well-tolerated and demonstrated antitumor activity in advanced mesothelioma and NF2-mutant cancers ^47^. In fact, directly targeting TEAD proteins to inhibit YAP1/TAZ activity may represent a more selective approach compared to blockade of Rho-GTPase signaling, as Rho-GTPase have several biological effector functions ^48^. To this end, various other small molecule TEAD inhibitors with promising pre-clinical activity have been disclosed in over the last years ^49^. Some of these inhibitors have progressed to the clinical stage^20,50^ and will hopefully shed light on the therapeutic potential of YAP1/TAZ inhibition in cancer.

## ACKNOWLEDGEMENTS

We thank Prof. Giannino Del Sal for kindly providing plasmids as a gift. Aurexel Life Sciences Ltd. (www.aurexel.com) is thanked for the editorial support funded by Bayer AG.

## AUTHOR CONTRIBUTIONS

Study Design, K. Graham and M. Lange; Data Analysis and Acquisition, K. Graham, B. Bader, S. Prechtl, J. Naujoks, R. Lesche, K. Brzezinka, B. Nicke, W. Bone, S. Golfier, S. Kaulfuss, C. Kopitz, H. Steuber, N. Braeuer, K. Nowak-Reppel, F. Zanconato, C. Stresemann, P. Steigemann, P. Lienau, J. Kuehnlenz, L. Potze, A. Montebaur, S. Pilari, P. Buchgraber, N. A. Font, T. Heinrich, L. Kuhnke, A.O. Walter, S. Blotta, M. Ocker, A. L. Eheim, D. Mumberg, K. Eis, S. Piccolo, M. Lange.; Bioinformatic Analyses, S. Hayat, A. Kamburov, A. Steffen, A. Schlicker; Study Supervision, K. Graham and M. Lange; Manuscript Preparation, K. Graham, P. Lienau, S. Prechtl, J. Naujoks, B. Nicke, M. Ocker, S. Piccolo, M. Lange

