## SUPPLEMENTARY DATA for "Novel YAP1/TAZ pathway inhibitors identified through phenotypic screening with potent anti-tumor activity via blockade of GGTase-I / Rho-GTPase signaling"

current affiliations:

[5] Boehringer Ingelheim Pharma GmbH & Co. KG, Biberach, Germany

[6] CELLIMA Scientific Services GmbH, Berlin

[7] Janssen Vaccines and Prevention, 2333 CB Leiden, The Netherlands

[8] Pfizer, Berlin, Germany

[9] BlueViolin GmbH, Berlin, Germany

[10] Institute of experimental medicine and systems biology, Uniklinik Aachen, Germany

[11] Owkin, Paris, France.

[12] BioNTech SE, 55131 Mainz, Germany

[13] Menarini Group, 50131 Firenze, Italy

[14] Tacalyx GmbH, 13353 Berlin, Germany

[15] F. Hoffmann-La Roche AG, CH-4070 Basel, Switzerland

[16] Adcendo ApS, 2200 Copenhagen, Denmark

### SUPPLEMENTARY FIGURES AND FIGURE LEGENDS

**Figure S1:**

Suppl. Figure S1, Graham et al.

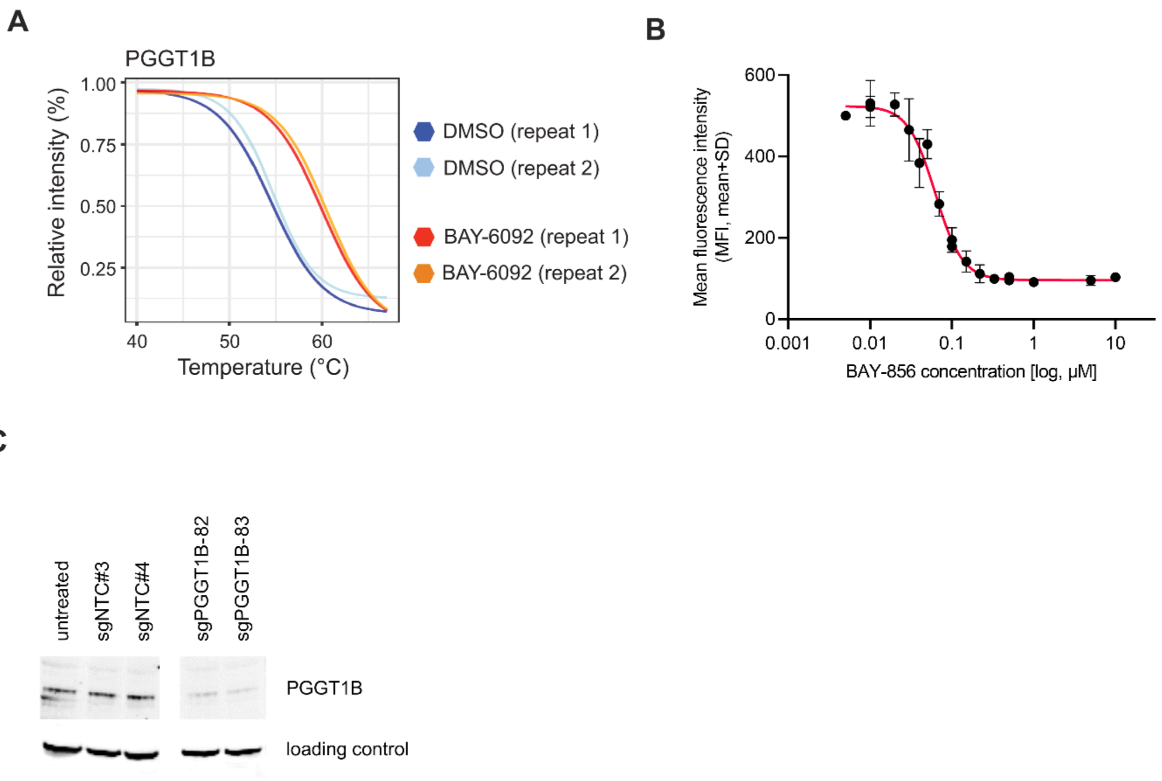

(A) BAY-6092 target engagement in MDA-MB-231 cell lysates treated with 10 μM BAY-6092 or vehicle control as determined by proteome-wide cellular thermal shift assay mass spectrometry (CETSA MS). Fold change values were normalized to a constant value of 1 by using the lowest temperature condition (+40°C) as reference

(B) $IC_{50}$ for BAY-856 was determined using a FACS-based luciferase expression assay, showing the fitted dose response curve of MDA-MB-231 TEAD-luc Cas9 cells treated for 24 hours with increasing concentrations of BAY-856. Cells were stained with firefly-luciferase antibody and analyzed by flow cytometry. Data is presented as mean fluorescence intensity (MFI) of luciferase signal, error bars present mean ± SD in biological triplicates.

(C) Knockdown of *PGGT1B* was confirmed by protein immunoblotting using anti-PGGT1B antibody. GAPDH was used as loading control.

**Figure S2:**

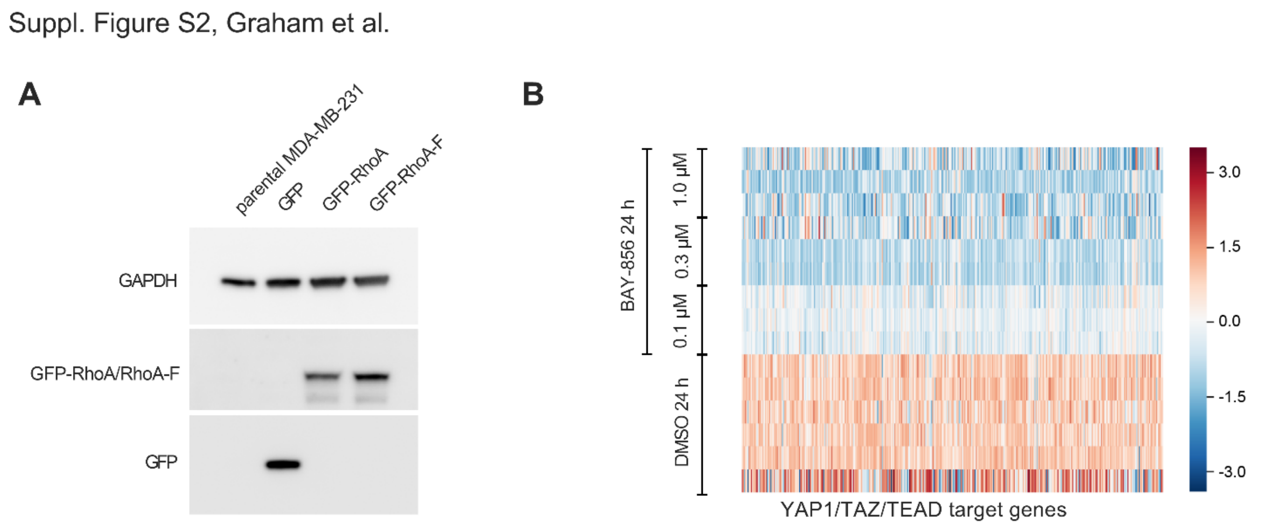

(A) Using protein immunoblotting, the protein expression of MDA-MB-231 cells transfected with either GFP-RhoA, GFP-RhoA-F or GFP was analyzed. GAPDH was used as loading control.

(B) Heatmap of YAP1/TAZ target gene mRNA expression changes by RNA-sequencing in MDA-MD-231 cells treated with DMSO or 0.1-1.0 µM BAY-856 for 24 hours

**Figure S3:**

Suppl. Figure S3, Graham et al

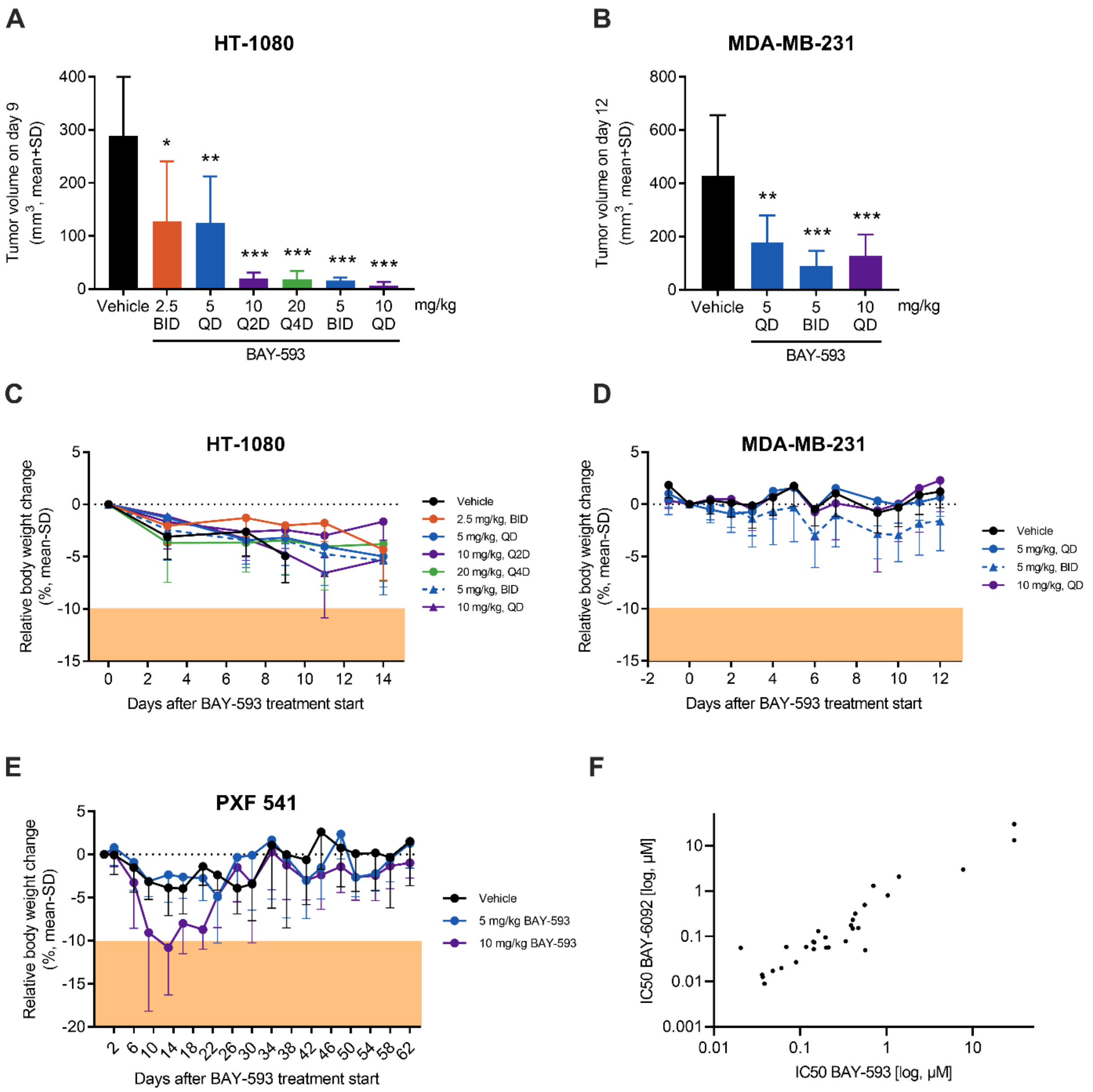

(A) HT-1080 tumor volumes were compared to the vehicle treatment on day 9. Variance was estimated separately for each group using weighted least squares: *, $p<0.05$; **, $p<0.01$; ***, $p<0.001$.

(B) MDA-MB-231 tumor volumes were compared to the vehicle treatment on day 12. ANOVA in log scale: **, $p<0.01$; ***, $p<0.001$.

(C) Relative body weight changes of mice with HT-1080 tumors. Mice were followed throughout the study.

(D) Relative body weight changes of mice with MDA-MB-231 tumors. Mice were followed throughout the study.

(E) Relative body weight changes of mice with PXF 541 tumors. Mice were followed throughout the study.

(F) Correlation of anti-proliferative activity (IC50 values) of BAY-6092 and BAY-593 based on cancer cell line screening.

### SUPPLEMENTARY TABLES

### Table S1: BAY-856 [1 μM] in kinase panel (Eurofins)

| Kinase | Kinase activity (%) | Kinase | Kinase activity (%) | Kinase | Kinase activity (%) | Kinase | Kinase activity (%) |
|---|---|---|---|---|---|---|---|
| Abl(h) | 102 | EphA2(h) | 90 | MEKK2(h) | 101 | PTK5(h) | 105 |
| ACK1(h) | 110 | EphA3(h) | 96 | MEKK3(h) | 92 | Pyk2(h) | 107 |
| ACTR2(h) | 97 | EphA4(h) | 99 | MELK(h) | 103 | Ret(h) | 108 |
| ALK(h) | 148 | EphA5(h) | 111 | Mer(h) | 98 | RIPK1(h) | 107 |
| ALK1(h) | 121 | EphA7(h) | 91 | Met(h) | 118 | RIPK2(h) | 98 |
| ALK2(h) | 108 | EphA8(h) | 112 | MINK(h) | 97 | ROCK-I(h) | 103 |
| ALK4(h) | 96 | EphB2(h) | 109 | MKK6(h) | 100 | ROCK-II(h) | 105 |
| ALK6(h) | 69 | EphB1(h) | 99 | MKK7β(h) | 104 | Ron(h) | 100 |
| Arg(h) | 124 | EphB3(h) | 98 | MLCK(h) | 92 | Ros(h) | 97 |
| AMPKα1(h) | 108 | EphB4(h) | 102 | MLK1(h) | 90 | Rse(h) | 148 |
| AMPKα2(h) | 107 | ErbB2(h) | 104 | MLK2(h) | 79 | Rsk1(h) | 108 |
| A-Raf(h) | 85 | ErbB4(h) | 114 | Mnk2(h) | 92 | Rsk2(h) | 100 |
| ARK5(h) | 105 | FAK(h) | 98 | MOK(h) | 78 | Rsk3(h) | 106 |
| ASK1(h) | 99 | Fer(h) | 108 | MRCKα(h) | 102 | Rsk4(h) | 113 |
| Aurora-A(h) | 123 | Fes(h) | 79 | MRCKβ(h) | 105 | SAPK2a(h) | 99 |
| Aurora-B(h) | 89 | FGFR1(h) | 99 | MSK1(h) | 100 | SAPK2b(h) | 92 |
| Aurora-C(h) | 94 | FGFR2(h) | 101 | MSK2(h) | 105 | SAPK3(h) | 106 |
| Axl(h) | 112 | FGFR3(h) | 97 | MSSK1(h) | 101 | SAPK4(h) | 94 |
| Blk(h) | 104 | FGFR4(h) | 108 | MST1(h) | 83 | SGK(h) | 105 |
| BMPR2(h) | 95 | Fgr(h) | 91 | MST2(h) | 100 | SGK2(h) | 102 |
| Bmx(h) | 105 | Flt1(h) | 112 | MST3(h) | 106 | SGK3(h) | 93 |
| BRK(h) | 94 | Flt3(h) | 96 | MST4(h) | 78 | SIK(h) | 87 |
| BrSK1(h) | 99 | Flt4(h) | 130 | mTOR(h) | 113 | SIK2(h) | 96 |
| BrSK2(h) | 109 | Fms(h) | 97 | mTOR/FKBP12(h) | 117 | SIK3(h) | 99 |
| BTK(h) | 103 | Fyn(h) | 109 | MuSK(h) | 105 | SLK(h) | 106 |
| B-Raf(h) | 101 | GCK(h) | 129 | MYLK2(h) | 95 | Snk(h) | 96 |
| CaMKI(h) | 98 | GCN2(h) | 90 | MYO3B(h) | 101 | SNRK(h) | 95 |
| CaMKIß(h) | 90 | GRK1(h) | 117 | NEK1(h) | 83 | SRPK1(h) | 105 |
| CaMKIγ(h) | 91 | GRK2(h) | 106 | NEK2(h) | 105 | SRPK2(h) | 111 |
| CaMKIIα(h) | 91 | GRK3(h) | 105 | NEK3(h) | 92 | STK25(h) | 107 |
| CaMKIIβ(h) | 102 | GRK5(h) | 93 | NEK6(h) | 104 | STK32A(h) | 94 |
| CaMKIIγ(h) | 95 | GRK6(h) | 96 | NEK7(h) | 107 | STK32B(h) | 96 |
| CaMKIδ(h) | 117 | GRK7(h) | 94 | NEK9(h) | 108 | STK32C(h) | 108 |
| CaMKIIδ(h) | 105 | GSK3α(h) | 90 | NIM1(h) | 101 | STK33(h) | 106 |
| CaMKIV(h) | 104 | GSK3β(h) | 101 | NEK11(h) | 91 | Syk(h) | 108 |
| CaMKK1(h) | 105 | Haspin(h) | 100 | NLK(h) | 91 | TAK1(h) | 100 |
| CaMKK2(h) | 116 | Hck(h) | 119 | NUAK2(h) | 97 | TAO1(h) | 95 |
| CDK1/cyclinB(h) | 103 | Hck(h) activated | 114 | p70S6K(h) | 104 | TAO2(h) | 100 |
| CDK2/cyclinA(h) | 87 | HIPK1(h) | 111 | PAK1(h) | 95 | TAO3(h) | 90 |
| CDK2/cyclinE(h) | 101 | HIPK2(h) | 105 | PAK2(h) | 104 | TBK1(h) | 110 |
| CDK3/cyclinE(h) | 110 | HIPK3(h) | 115 | PAK4(h) | 93 | Tec(h) activated | 101 |
| CDK4/cyclinD3(h) | 113 | HIPK4(h) | 120 | PAK3(h) | 98 | TGFBR1(h) | 97 |
| CDK5/p25(h) | 100 | HPK1(h) | 102 | PAK5(h) | 90 | Tie2 (h) | 98 |
| CDK5/p35(h) | 102 | HRI(h) | 94 | PAK6(h) | 92 | TLK1(h) | 90 |
| CDK6/cyclinD3(h) | 99 | ICK(h) | 101 | PAR-1Bα(h) | 90 | TLK2(h) | 107 |
| CDK7/cyclinH/MAT1(h) | 91 | IGF-1R(h) | 104 | PASK(h) | 113 | TNIK(h) | 112 |
| CDK9/cyclin T1(h) | 104 | IGF-1R(h), activated | 108 | PEK(h) | 113 | TrkA(h) | 101 |
| CDK12/cyclinK(h) | 115 | IKKα(h) | 99 | PDGFRα(h) | 100 | TrkB(h) | 121 |
| CDK18/cyclinY(h) | 137 | IKKβ(h) | 100 | PDGFRβ(h) | 85 | TrkC(h) | 101 |
| CDKL1(h) | 94 | IKKε(h) | 102 | PDHK4(h) | 94 | TSSK1(h) | 98 |
| ChaK1(h) | 101 | IR(h) | 138 | PDK1(h) | 83 | TSSK2(h) | 109 |
| CHK1(h) | 95 | IR(h), activated | 100 | PhKγ1(h) | 95 | TSSK3(h) | 103 |
| CHK2(h) | 97 | IRE1(h) | 110 | PhKγ2(h) | 106 | TSSK4(h) | 106 |
| CK1γ1(h) | 96 | IRR(h) | 100 | Pim-1(h) | 113 | TTBK1(h) | 95 |
| CK1γ2(h) | 97 | IRAK1(h) | 97 | Pim-2(h) | 92 | TTBK2(h) | 95 |
| CK1γ3(h) | 122 | IRAK4(h) | 91 | Pim-3(h) | 87 | TTK(h) | 97 |
| CK1δ(h) | 100 | Itk(h) | 102 | PKA(h) | 98 | Txk(h) | 96 |
| CK2(h) | 95 | JAK1(h) | 103 | PKAcβ(h) | 107 | TYK2(h) | 101 |
| CK2α1(h) | 83 | JAK2(h) | 114 | PKBα(h) | 105 | ULK1(h) | 99 |
| CK2α2(h) | 84 | JAK3(h) | 108 | PKBβ(h) | 109 | ULK2(h) | 92 |
| CLIK1(h) | 94 | JNK1α1(h) | 111 | PKBγ(h) | 92 | ULK3(h) | 89 |
| CLK1(h) | 101 | JNK2α2(h) | 99 | PKCα(h) | 98 | Wee1(h) | 96 |

| | | | | | | | |
|---|---|---|---|---|---|---|---|
| CLK2(h) | 102 | JNK3(h) | 115 | PKCβI(h) | 94 | Wee1B(h) | 95 |
| CLK3(h) | 98 | KDR(h) | 90 | PKCβII(h) | 90 | WNK2(h) | 99 |
| CLK4(h) | 84 | Lck(h) | 116 | PKCγ(h) | 105 | WNK3(h) | 95 |
| cKit(h) | 98 | Lck(h) activated | 103 | PKCδ(h) | 113 | VRK2(h) | 104 |
| CSK(h) | 114 | LIMK1(h) | 105 | PKCε(h) | 92 | Yes(h) | 98 |
| c-RAF(h) | 102 | LKB1(h) | 104 | PKCη(h) | 95 | ZAK(h) | 100 |
| cSRC(h) | 91 | LOK(h) | 116 | PKCι(h) | 87 | ZAP-70(h) | 103 |
| DAPK1(h) | 93 | Lyn(h) | 108 | PKCμ(h) | 106 | ZIPK(h) | 112 |
| DAPK2(h) | 101 | LRRK2(h) | 112 | PKCθ(h) | 106 | ATM(h) | 102 |
| DCAMKL2(h) | 88 | LTK(h) | 93 | PKCζ(h) | 113 | ATR/ATRIP(h) | 109 |
| DCAMKL3(h) | 92 | MAK(h) | 93 | PKD2(h) | 94 | DNA-PK(h) | 97 |
| DDR1(h) | 121 | MAPK1(h) | 92 | PKD3(h) | 103 | PI3 Kinase (p110b/p85a)(h) | 100 |
| DDR2(h) | 107 | MAPK2(h) | 113 | PKG1α(h) | 106 | PI3 Kinase (p120g)(h) | 99 |
| DMPK(h) | 114 | MAP4K3(h) | 87 | PKG1β(h) | 96 | PI3 Kinase (p110d/p85a)(h) | 101 |
| DRAK1(h) | 105 | MAP4K4(h) | 100 | PKR(h) | 95 | PI3 Kinase (p110a/p85a)(h) | 98 |
| DRAK2(h) | 90 | MAP4K5(h) | 110 | Plk1(h) | 80 | PI3KC2a(h) | 102 |
| DYRK1A(h) | 107 | MAPKAP-K2(h) | 100 | Plk3(h) | 99 | PI3KC2g(h) | 103 |
| DYRK1B(h) | 101 | MAPKAP-K3(h) | 105 | PRAK(h) | 86 | PIP4K2a(h) | 102 |
| DYRK2(h) | 102 | MEK1(h) | 90 | PRKG2(h) | 95 | PIP5K1a(h) | 102 |
| DYRK3(h) | 94 | MEK2(h) | 96 | PRK1(h) | 106 | PIP5K1g(h) | 98 |
| eEF-2K(h) | 102 | MARK1(h) | 93 | PRK2(h) | 101 | | |
| EGFR(h) | 102 | MARK3(h) | 104 | PrKX(h) | 89 | | |
| EphA1(h) | 100 | MARK4(h) | 83 | PRP4(h) | 101 | | |

### Table S2: CRISPR primers

| sgRNAs | Primer ID |
|---|---|
| Non-targeting control #1 | VSGC10215 |
| Non-targeting control #2 | VSGC10216 |
| Non-targeting control #3 | VSGC10217 |
| WWTR1 | VSGHSM_26670159, VSGHSM_26670158, VSGHSM_26670157 |
| **NGS sequencing primers** | **Primer sequence** |
| Forward primer (FwdU6-1) | 5' CAAGGCTGTTAGAGAGATAATTGG 3' |
| Reverse primer (Rev-1) | 5' CGACAACAACGCAGAAATTTTGAAT 3' |
| P7-NFG16 | CAAGCAGAAGACGGCATACGAGATGCATGTATGTCTGTTGCTATTATGTCTACT |
| P5-NFG16-IndA | AATGATACGGCGACCACCGAGATCTACACTACGACATGGACTATCATATGCTTACCGTAACTTGAA |
| P5-NFG16-IndB | AATGATACGGCGACCACCGAGATCTACACCTGATGATGGACTATCATATGCTTACCGTAACTTGAA |
| P5-NFG16-IndC | AATGATACGGCGACCACCGAGATCTACACGCATCAATGGACTATCATATGCTTACCGTAACTTGAA |
| P5-NFG16-IndD | AATGATACGGCGACCACCGAGATCTACACAGTCGTATGGACTATCATATGCTTACCGTAACTTGAA |
| P5-NFG16-IndE | AATGATACGGCGACCACCGAGATCTACACTCGCATATGGACTATCATATGCTTACCGTAACTTGAA |
| P5-NFG16-IndF | AATGATACGGCGACCACCGAGATCTACACCATAGCATGGACTATCATATGCTTACCGTAACTTGAA |
| P5-NFG16-IndG | AATGATACGGCGACCACCGAGATCTACACAGCGTAATGGACTATCATATGCTTACCGTAACTTGAA |
| P5-NFG16-IndH | AATGATACGGCGACCACCGAGATCTACACGTATCGATGGACTATCATATGCTTACCGTAACTTGAA |
| P5-NFG16-IndP | AATGATACGGCGACCACCGAGATCTACACATCAGCATGGACTATCATATGCTTACCGTAACTTGAA |
| **Sequencing Primer** | **Primer sequence** |
| FSeq-gRNA | GGCTTTATATATCTTGTGGAAAGGACGAAACACCG |

To construct the sequencing library, the primer P7-NFG16 was internally modified by reducing the primer's length. The provided P5-NFG16 primer was extended by 6 bases (indicated red) to generate distinguishable index primers. The index sequence was introduced directly after the Illumina P5 adapter sequence. Primers contain required Illumina P5 and P7 sequences for next generation sequencing on MiSeq system (Illumina).

### Table S3: Cell proliferation screen of 250 human cancer cell lines treated with BAY-6092

| Cell line | Activity Area | IC50 [M] |
|---|---|---|
| HT1080 | 7,82 | 3,34E-09 |
| BHT101 | 6,71 | 4,92E-09 |
| MDAMB231 | 6,36 | 4,09E-09 |
| 769P | 6,15 | 1,16E-08 |
| CAKI1 | 5,81 | 6,07E-09 |
| HUPT4 | 5,74 | 9,95E-09 |
| SW982 | 5,63 | 1,83E-09 |
| DU145 | 5,59 | 1,96E-08 |
| SCC9 | 5,37 | 6,48E-09 |
| BPH1 | 5,37 | 2,04E-08 |
| H4 | 5,36 | 5,88E-09 |
| SNB19 | 5,33 | 1,08E-08 |
| T24 | 5,24 | 2,08E-09 |
| OE21 | 5,13 | 2,56E-08 |
| VAESBJ | 5,03 | 1,07E-08 |
| DETROIT562 | 5,01 | 1,05E-08 |
| NCIH292 | 4,9 | 1,17E-08 |
| ACHN | 4,85 | 1,46E-08 |
| 786O | 4,85 | 3,99E-08 |
| HUTU80 | 4,78 | 1,12E-07 |
| CAL62 | 4,73 | 4,21E-09 |
| NCIH460 | 4,61 | 1,20E-08 |
| G292CLONEA141B1 | 4,6 | 1,65E-07 |
| PA1 | 4,55 | 1,66E-08 |
| PANC1 | 4,55 | 2,06E-08 |
| BM1604 | 4,5 | 7,65E-08 |
| SW954 | 4,48 | 1,03E-08 |
| T98G | 4,47 | 1,43E-08 |
| HPAFII | 4,47 | 1,21E-07 |
| A172 | 4,4 | 1,95E-08 |
| CAL27 | 4,36 | 8,69E-09 |
| PC3PRO | 4,35 | 4,41E-08 |
| A431 | 4,2 | 9,63E-09 |
| UMUC3 | 4,08 | 1,03E-08 |
| CORL105 | 4,07 | 1,14E-07 |
| OCUG1 | 4,02 | 1,10E-08 |
| HS766T | 4,01 | 3,00E-05 |
| RPMI7951 | 4 | 9,85E-09 |
| HLF | 4 | 1,86E-08 |
| PSN1 | 4 | 2,09E-08 |
| SW684 | 3,99 | 3,00E-05 |
| HLE | 3,97 | 1,46E-08 |
| RKO | 3,96 | 9,77E-09 |
| SKLMS1 | 3,94 | 7,99E-09 |
| HELA | 3,93 | 1,17E-07 |
| SW872 | 3,91 | 1,50E-07 |
| SW13 | 3,88 | 1,29E-08 |
| SW1353 | 3,83 | 3,33E-08 |
| HS746T | 3,79 | 9,72E-09 |
| CALU1 | 3,79 | 1,02E-08 |
| 647V | 3,78 | 5,96E-08 |
| 5637 | 3,77 | 6,12E-08 |
| SW900 | 3,72 | 3,00E-05 |
| A375 | 3,7 | 5,82E-09 |
| DOTC24510 | 3,66 | 7,53E-07 |
| FADU | 3,59 | 6,16E-07 |
| OE33 | 3,53 | 1,03E-06 |
| G402 | 3,51 | 2,59E-08 |
| A427 | 3,47 | 3,94E-07 |
| RL952 | 3,47 | 3,00E-05 |
| CAKI2 | 3,44 | 2,07E-08 |
| HS294T | 3,44 | 2,66E-08 |
| CHL1 | 3,44 | 1,83E-07 |
| U87MG | 3,38 | 3,00E-05 |

| Cell line | Activity Area | IC50 [M] |
|---|---|---|
| MDAMB468 | 2,46 | 3,00E-05 |
| MESSA | 2,44 | 3,00E-05 |
| D341MED | 2,44 | 3,00E-05 |
| MEWO | 2,43 | 3,00E-05 |
| SCC25 | 2,4 | 2,05E-07 |
| SNU1 | 2,39 | 3,00E-05 |
| SKUT1 | 2,38 | 3,00E-05 |
| MG63 | 2,36 | 3,00E-05 |
| SNU5 | 2,36 | 3,00E-05 |
| HT3 | 2,35 | 1,45E-05 |
| NCIH747 | 2,34 | 2,87E-05 |
| SJCRH30 | 2,34 | 3,00E-05 |
| SW962 | 2,34 | 3,00E-05 |
| BXPC3 | 2,33 | 3,00E-05 |
| DMS53 | 2,33 | 3,00E-05 |
| SJSA1 | 2,32 | 3,00E-05 |
| SW1463 | 2,31 | 3,00E-05 |
| HCT116 | 2,27 | 3,00E-05 |
| HCT15 | 2,27 | 3,00E-05 |
| SNUC2B | 2,26 | 3,00E-05 |
| ME180 | 2,25 | 3,00E-05 |
| G401 | 2,24 | 2,29E-06 |
| NCIH596 | 2,22 | 3,00E-05 |
| HUCCT1 | 2,21 | 3,00E-05 |
| KATOIII | 2,21 | 3,00E-05 |
| SW1783 | 2,19 | 3,00E-05 |
| SKMEL28 | 2,18 | 3,00E-05 |
| SW1417 | 2,17 | 3,00E-05 |
| CALU6 | 2,16 | 3,00E-05 |
| HS729 | 2,14 | 3,00E-05 |
| MCIXC | 2,12 | 3,00E-05 |
| AN3CA | 2,1 | 3,00E-05 |
| SNU16 | 2,1 | 3,00E-05 |
| SW837 | 2,1 | 3,00E-05 |
| JAR | 2,07 | 7,97E-06 |
| SH4 | 2,07 | 3,00E-05 |
| OE19 | 2,06 | 3,00E-05 |
| CASKI | 2,06 | 3,00E-05 |
| KLE | 2,06 | 3,00E-05 |
| M059J | 2,05 | 3,00E-05 |
| SKMEL3 | 2,05 | 3,00E-05 |
| U2OS | 2,05 | 3,00E-05 |
| TCCSUP | 2,03 | 3,00E-05 |
| YAPC | 2,01 | 3,00E-05 |
| CCFSTTG1 | 1,99 | 3,00E-05 |
| HMCB | 1,99 | 3,00E-05 |
| WIDR | 1,97 | 3,00E-05 |
| HT1197 | 1,9 | 3,00E-05 |
| HS852T | 1,89 | 3,00E-05 |
| DMS114 | 1,89 | 3,00E-05 |
| C33A | 1,88 | 3,00E-05 |
| MCF7 | 1,87 | 3,00E-05 |
| ZR751 | 1,85 | 3,00E-05 |
| C4II | 1,84 | 3,00E-05 |
| SW480 | 1,84 | 3,00E-05 |
| A549 | 1,82 | 3,00E-05 |
| DOHH2 | 1,78 | 1,06E-05 |
| WM2664 | 1,74 | 3,00E-05 |
| CAMA1 | 1,73 | 3,00E-05 |
| DLD1 | 1,72 | 1,42E-05 |
| C4I | 1,72 | 3,00E-05 |
| HS695T | 1,66 | 3,00E-05 |
| BEWO | 1,61 | 3,00E-05 |
| G361 | 1,61 | 3,00E-05 |

| | | |
|---|---|---|
| NCIH661 | 3,37 | 3,00E-05 |
| HOS | 3,36 | 4,77E-08 |
| J82 | 3,35 | 3,00E-05 |
| A388 | 3,34 | 4,00E-08 |
| HCT8 | 3,32 | 2,15E-08 |
| HEC1A | 3,31 | 8,47E-08 |
| HS688AT | 3,31 | 3,00E-05 |
| SKMES1 | 3,3 | 3,00E-05 |
| DMS273 | 3,29 | 7,72E-08 |
| D283MED | 3,25 | 3,73E-07 |
| DKMG | 3,23 | 3,00E-05 |
| RDSOF | 3,21 | 3,00E-05 |
| DBTRG05MG | 3,2 | 3,87E-08 |
| C32TG | 3,18 | 5,91E-08 |
| SKOV3 | 3,18 | 3,00E-05 |
| HS895T | 3,17 | 3,00E-05 |
| COLO205 | 3,16 | 4,29E-06 |
| BFTC905 | 3,14 | 5,65E-08 |
| 639V | 3,14 | 6,51E-08 |
| A204 | 3,12 | 3,00E-05 |
| Y79 | 3,11 | 3,00E-05 |
| COLO829 | 3,08 | 3,00E-05 |
| SKNFI | 3,08 | 3,00E-05 |
| AGS | 3,05 | 3,98E-07 |
| 22RV1 | 3,05 | 3,79E-06 |
| CGTHW1 | 3,04 | 3,00E-05 |
| KHOS240S | 3,03 | 6,00E-08 |
| A7 | 3,03 | 2,19E-07 |
| SCABER | 3,02 | 4,91E-08 |
| MDAMB415 | 3,01 | 3,00E-05 |
| SCC4 | 3,01 | 3,00E-05 |
| SNU423 | 3 | 4,95E-08 |
| TE381T | 3 | 7,37E-08 |
| CAOV3 | 2,94 | 2,74E-07 |
| U138MG | 2,91 | 3,00E-05 |
| SW579 | 2,9 | 1,52E-05 |
| MIAPACA2 | 2,89 | 1,16E-07 |
| SW1088 | 2,87 | 1,16E-07 |
| A498 | 2,86 | 9,89E-08 |
| RKOAS451 | 2,85 | 3,00E-05 |
| HT29 | 2,84 | 3,00E-05 |
| NTERA2CLD1 | 2,8 | 9,41E-08 |
| DAOY | 2,78 | 2,19E-06 |
| A2058 | 2,75 | 3,00E-05 |
| A673 | 2,74 | 3,00E-05 |
| SIHA | 2,73 | 3,00E-05 |
| PFSK1 | 2,72 | 2,88E-05 |
| NCIH446 | 2,71 | 3,00E-05 |
| LS513 | 2,7 | 1,28E-05 |
| OVCAR3 | 2,69 | 1,64E-05 |
| SKNEP1 | 2,62 | 2,37E-06 |
| A101D | 2,62 | 3,00E-05 |
| SUDHL4 | 2,61 | 4,90E-08 |
| COLO320HSR | 2,61 | 3,00E-05 |
| HS683 | 2,61 | 3,00E-05 |
| CHP212 | 2,6 | 3,00E-05 |
| JURKAT | 2,54 | 3,00E-05 |
| HUH6CLONE5 | 2,52 | 7,09E-08 |
| HS578T | 2,52 | 3,00E-05 |
| A704 | 2,49 | 3,00E-05 |
| RKOE6 | 2,47 | 1,06E-06 |

| | | |
|---|---|---|
| LS123 | 1,6 | 3,00E-05 |
| SAOS2 | 1,59 | 3,00E-05 |
| ASPC1 | 1,55 | 3,00E-05 |
| SW403 | 1,54 | 3,00E-05 |
| NCIH441 | 1,54 | 3,00E-05 |
| HS821T | 1,47 | 3,00E-05 |
| HS936TC1 | 1,46 | 3,00E-05 |
| MDAMB436 | 1,4 | 3,00E-05 |
| HEPG2 | 1,39 | 3,00E-05 |
| NCIH510A | 1,38 | 3,00E-05 |
| NCIH520 | 1,36 | 3,00E-05 |
| MALME3M | 1,35 | 3,00E-05 |
| COLO201 | 1,33 | 3,00E-05 |
| A253 | 1,3 | 3,00E-05 |
| MT3 | 1,29 | 3,00E-05 |
| SW620 | 1,29 | 3,00E-05 |
| CAPAN1 | 1,28 | 3,00E-05 |
| SU8686 | 1,24 | 3,00E-05 |
| U266B1 | 1,24 | 3,00E-05 |
| AU565 | 1,2 | 2,36E-05 |
| SW48 | 1,2 | 3,00E-05 |
| KPL1 | 1,19 | 1,74E-05 |
| HS888SK | 1,18 | 3,00E-05 |
| U118MG | 1,14 | 3,00E-05 |
| NALM6 | 1,13 | 3,00E-05 |
| C32 | 1,11 | 3,00E-05 |
| LNCAP | 1,07 | 2,34E-05 |
| BT474 | 1,07 | 3,00E-05 |
| LS1034 | 1,04 | 3,00E-05 |
| SKNAS | 1,02 | 3,00E-05 |
| SKBR3 | 1 | 3,00E-05 |
| DAUDI | 0,97 | 3,00E-05 |
| HS229T | 0,97 | 3,00E-05 |
| SKPNDW | 0,96 | 3,00E-05 |
| IM9 | 0,92 | 3,00E-05 |
| LS174T | 0,92 | 3,00E-05 |
| MS751 | 0,91 | 3,00E-05 |
| SHP77 | 0,9 | 3,00E-05 |
| BT20 | 0,87 | 3,00E-05 |
| CORL23 | 0,87 | 3,00E-05 |
| COLO320DM | 0,83 | 3,00E-05 |
| JEG3 | 0,8 | 2,83E-05 |
| MDAMB453 | 0,76 | 3,00E-05 |
| CAPAN2 | 0,74 | 3,00E-05 |
| TTTHY | 0,72 | 3,00E-05 |
| T47D | 0,66 | 3,00E-05 |
| BE2C | 0,65 | 2,86E-05 |
| LS411N | 0,58 | 3,00E-05 |
| CHAGOK1 | 0,53 | 3,00E-05 |
| NCIH508 | 0,44 | 3,00E-05 |
| NCIH295R | 0,43 | 3,00E-05 |
| HS934T | 0,42 | 3,00E-05 |
| EFM19 | 0,32 | 3,00E-05 |
| KG1 | 0,3 | 3,00E-05 |
| NCIH69 | 0,3 | 3,00E-05 |
| SW948 | 0,24 | 3,00E-05 |
| TE125T | 0,18 | 3,00E-05 |
| SKNDZ | 0,17 | 3,00E-05 |
| SKMEL1 | 0,16 | 3,00E-05 |
| BT549 | 0 | 3,00E-05 |
| HT1376 | 0 | 3,00E-05 |

### Table S4: Key Resources Table

| REAGENT or RESOURCE | SOURCE | IDENTIFIER |
|---|---|---|
| Antibodies | | |
| YAP (D8H1X) XP® Rabbit mAb Alexa Fluor® 488 Conjugate | Cell Signaling (CST) | #14729 |
| YAP (D8H1X) XP® Rabbit mAb | Cell Signaling (CST) | #14074 |
| Hoechst 33342 | Thermo Fisher Scientific | H3570 |
| Recombinant Anti-Firefly Luciferase antibody [EPR17790] | Abcam | ab185924 |
| Alexa Fluor 488 Goat anti-Rabbit IgG secondary antibody | Thermo Fisher Scientific | A-11008 |
| WWTR1 (TAZ) Rabbit polyclonal Ab | Atlas Antibodies | HPA007415 |
| Alexa Fluor™ 488 Phalloidin conjugate | Invitrogen | A12379 |
| α-Tubulin antibody | CST | #2144 |
| GAPDH (14C10) Rabbit mAb | CST | #2118 |
| Calreticulin antibody | CST | #2891 |
| CYR61 (D4H5D) XP® Rabbit mAb | CST | #14479 |
| Recombinant Anti-PD-L1 antibody [EPR20529] | Abcam | ab213480 |
| RhoA (67B9) Rabbit mAb | CST | #2117 |
| RhoC (D40E4) Rabbit mAb | CST | #3430 |
| Rac1/Cdc42 antibody | CST | #4651 |
| Cdc42 antibody | CST | #2462 |
| Anti-Glyceraldehyde-3-Phosphate Dehydrogenase antibody, clone 6C5 | Millipore | MAB374 |
| GFP antibody (FL) | Santa Cruz Biotechnology | sc-8334 |
| anti-mouse HRP | Abcam | ab97023 |
| anti-rabbit HRP | Abcam | ab205718 |
| Chemicals, Peptides, and Recombinant Proteins | | |
| BAY-856 / BAY-6092 / BAY-593 | Bayer AG | |
| C3 exoenzyme | Tocris Bioscience | |
| Cerivastatin | Tocris Bioscience | |
| GGTI-298 | Tocris Bioscience | |
| GGTI-2133 | Tocris Bioscience | |
| Lonafarnib | Tocris Bioscience | |
| Critical Commercial Assays | | |
| DualGlo™luciferase assay system detection kit | Promega | #E2920, #E2940 |
| Dual-Glo™ Stop&Glo Luciferase detection solution | | |
| CRISPRuTest™ Functional Cas9 Activity Assay kit | Cellecta | |
| Dual-Glo Luciferase Assay System | Promega | |
| Fixation/Permeabilization Solution kit | BD Biosciences | |
| GeneRead DNA FFPE kit | Qiagen | |
| Subcellular Protein Fractionation kit | Thermo Fischer Scientific | #7884 |
| RNAeasy kit | Qiagen | #74181 |
| LunaScript® RT SuperMix kit | New England Biolabs | #E3010 |
| Taqman Fast Advanced Master Mix | Thermo Fisher Scientific | #4444965 |
| TruSeq Stranded mRNA Library Prep kit | Illumina | RS-122-2103 |
| HiSeq SR Cluster Kit v4 cBOT | Illumina | GD-401-4001 |
| HiSeq SBS Kit v4 | Illumina | FC-401-4002 |
| CellTiter-Glo (CTG) solution | Promega | #G755B |
| Retrieval solution, pH 6 | Agilent Technologies | S2031 |
| EnVision+ System-HRP | Agilent Technologies / Dako | #K4010 |
| Fluoroshield™ with DAPI | Sigma-Aldrich | F6057 |

| | | |
|---|---|---|
| Pierce™ ECL Plus Western Blotting Substrate | Pierce | #32132 |
| Deposited Data | | |
| Protein Geranylgeranyltransferase type-I Complexed with Geranylgeranyl Diphosphate | Taylor et al., 2003 | PDB entry 1N4P |
| Gencode human reference genome (assembly hg19, GRCh37.p13) | | https://www.gencodegenes.org/human/release_19.html |
| Experimental Models: Cell Lines | | |
| 769P | AATC / DSMZ | - |
| 786O | AATC / DSMZ | - |
| A-2058 | AATC / DSMZ | - |
| CAL27 | AATC / DSMZ | - |
| CFPAC-1 | AATC / DSMZ | - |
| CHL-1 | AATC / DSMZ | - |
| HEL92.1.7 | AATC / DSMZ | - |
| H1373 | AATC / DSMZ | - |
| H358 | AATC / DSMZ | - |
| HT-1080 | AATC / DSMZ | - |
| JJN-3 | AATC / DSMZ | - |
| MDA-MB-231 | AATC / DSMZ | - |
| MOLM-16 | AATC / DSMZ | - |
| SCC9 | AATC / DSMZ | - |
| SKMEL-3 | AATC / DSMZ | - |
| T24 | AATC / DSMZ | - |
| T98G | AATC / DSMZ | - |
| Experimental Models: Organisms/Strains | | |
| Mouse: NMRI nu/nu mice | Janvier | |
| Mouse: nu/nu mice | Charles River | |
| Oligonucleotides | | |
| 150k 3-Module Human Genome-Wide CRISPR Library | Cellecta | |
| ACTB Taqman probe | Thermo Fisher Scientific | 4333762 |
| ANKRD1 Taqman probe | Thermo Fisher Scientific | Hs00923599_m1 |
| CCNA2 Taqman probe | Thermo Fisher Scientific | Hs00996788_m1 |
| CD274 Taqman probe | Thermo Fisher Scientific | Hs00204257_m1 |
| CDC6 Taqman probe | Thermo Fisher Scientific | Hs00154374_m1 |
| CDCA5 Taqman probe | Thermo Fisher Scientific | Hs00293564_m1 |
| CENPF Taqman probe | Thermo Fisher Scientific | Hs01118845_m1 |
| CTGF Taqman probe | Thermo Fisher Scientific | Hs01026927_g1 |
| CYR61 Taqman probe | Thermo Fisher Scientific | Hs0098500_g1 |
| DKK1 Taqman probe | Thermo Fisher Scientific | Hs00183740_m1 |
| ETS1 Taqman probe | Thermo Fisher Scientific | Hs00428293_m1 |
| FOSL1 Taqman probe | Thermo Fisher Scientific | Hs04187685_m1 |
| GAPDH Taqman probe | Thermo Fisher Scientific | 4333764 |
| GINS1 Taqman probe | Thermo Fisher Scientific | Hs00221421_m1 |
| KIF23 Taqman probe | Thermo Fisher Scientific | Hs00370852_m1 |
| MCM3 Taqman probe | Thermo Fisher Scientific | Hs00172459_m1 |
| MCM7 Taqman probe | Thermo Fisher Scientific | Hs00428518_m1 |
| MYBL1 Taqman probe | Thermo Fisher Scientific | Hs00277143_m1 |
| PPIA Taqman probe | Thermo Fisher Scientific | 4333763 |
| Recombinant DNA | | |
| Luc2P firefly luciferase reporter | Promega | pGL4.27 vector |
| Thymidine kinase (TK)-Renilla reporter | Promega | pGL4.74 vector |
| pLv5-Cas9-Neo | Sigma-Aldrich | |
| CMV-TagGFP2-KRAS$^{G12C}$-IRES-Neo | | |
| GFP-RhoA | Sorrentino et al., 2014 | |
| GFP-RhoA-F | Sorrentino et al., 2014 | |
| TEAD reporter (8xGTIIC-lux) | Dupont et al., 2011 | |
| Software and Algorithms | | |

| | | |
|---|---|---|
| MetaXpress$^{TM}$ image analysis software v5.1.0.41/v6.2.3 | Molecular Devices | |
| Sequest HT algorithm, ProteomeDiscoverer 2.1 software package | | |
| Genedata Assay Analyzer software v13.0 | Genedata | |
| Bowtie short read aligner open-source software v2.2.6 | | |
| XDS program package. | | |
| PHASER | Mccoy et al., 2007 | |
| COOT Model building | Emsley et al., 2010 | |
| REFMAC model refinement | Murshudov et al., 1997 | - |
| MathIQ based software (AIM). | | |
| STAR aligner alignment tool v2.4.2a | | |
| FeatureCounts v1.4.6-p2 from the Subread package | | |
| Bioconductor package DESeq2 | | https://bioconductor.org/packages/release/bioc/html/DESeq2.html |
| GSEA application v2.2 | | https://www.gsea-msigdb.org/gsea/index.jsp |
| Genedata Screener commercial software package | Genedata | |
| GraphPad Prism v8.0.2 | GraphPad Software Inc. | https://www.graphpad.com/scientific-software/prism/ |
| R language v3.6.3 | | https://www.r-project.org/ |
| Boruta v6.0.0 (R package) | | http://www.jstatsoft.org/v36/i11/ |
| the pamr package v1.55 | | |
| Other | | |
| OncoPanel™ Cell Proliferation screening platform | Eurofins | |
